# Mediator-RNA Polymerase II Interactions Critical for Transcriptional Activation Are Mediated by the N-terminal Half of MED14 and C-terminal Domain of RPB1

**DOI:** 10.1101/2022.04.27.489650

**Authors:** Yasemin Baris, Javaid Jabbar, Yasemin Yozgat, Ege Cigirgan, Sadik Bay, Merve Erden Tucer, Volkan Aslan, Melike Dinccelik-Aslan, Murat Alper Cevher

## Abstract

Mediator is a large and evolutionarily conserved coactivator complex essential for RNA polymerase II (Pol II)-mediated gene regulation at multiple steps of the transcription process, including preinitiation complex (PIC) assembly and function. Here, we used the MultiBac baculovirus expression system to generate recombinant human core Mediator subcomplexes and subsequent biochemical approaches to dissect the mechanism by which Mediator facilitates direct recruitment of Pol II to core promoters. Our results highlight a pivotal role in this process for the N-terminal half (NTD) of the MED14 subunit. We show that a reconstituted 15-subunit human core Mediator complex that contains only the MED14-NTD is fully functional in facilitating both basal and activated (p53) transcription. This complex directly interacts with the C-terminal domain (CTD) of the RPB1 subunit of Pol II (RPB1 CTD) and is required for recruiting Pol II to core promoters. Moreover, recombinant RPB1 can completely reverse the human core Mediator-Pol II interaction. Notably, the human MED14-NTD region has secondary structure conservation with *Schizosaccharomyces pombe*. In addition, reanalysis of published cryo-EM structures of yeast Mediator-Pol II complexes strongly supports our conclusion. Thus, our analyses provide critical new insights into how Mediator binds to Pol II and recruits it to the promoters to facilitate transcription.

## INTRODUCTION

In the paradigmatic case, transcriptional activation of protein-encoding genes by Pol II entails several steps^1–4^. In an initial stage, enhancer-bound activators recruit a series of chromatin remodeling and modifying cofactors that together covalently modify histones for recognition by other factors and evict nucleosomes to expose DNA regulatory elements^1–4^. PIC assembly is regulated by enhancer-bound activators that recruit additional coactivators, of which the Mediator is perhaps the single most critical. Upon binding to activators, Mediator is believed to facilitate recruitment and function of not only Pol II and general transcription factors (GTFs) but also various elongation factors that include the super elongation complex^1–4^.

Despite its large size (30 subunits and 2 MDa in humans), the Mediator complex displays a modular organization consisting of the head, middle, kinase, and tail modules, making it relatively amenable to a structure-function dissection^5^. A subset of subunits from the head and middle modules constitute the functional core Mediator that executes its effector functions through direct interactions with Pol II and several GTFs and shows limited activator (e.g., p53) interactions^1–4,6,7^. The tail and kinase modules are variably associated with the core Mediator and appear to have primary functions in regulating enhancer-promoter interactions (kinase module) and in Mediator recruitment through interactions with enhancer-bound activators (tail subunits)^1–4^.

One of the biggest challenges in the transcription field relates to our understanding of the detailed mechanism by which the Mediator binds Pol II and recruits it to promoters to facilitate transcription. As many of the Mediator subunits are essential, knockout and even deletion mutations of these subunits are not possible. Thus, only limited information could be gathered so far from studies that have employed cross-linking experiments, structure analysis, and genetic approaches to assign specific roles or interacting partners to the Mediator. Therefore, the nature and functions of Mediator subunits critical for the facilitation of PIC assembly remain unaddressed.

Several lines of evidence also previously implicated MED14 as a critical component for Mediator function in yeast. For instance, initial genetic screening of the Pol II-interacting proteins in yeast also identified MED14^8,9^. In addition, an N-terminal MED14 truncation extending to amino acid 473 (within the N-terminal half of MED14 in repeat motif two) was also lethal^9–11^.

All functional human Mediator isolates have been found to contain MED14. Notably, MED14 is a component of the TRAP complex, the earliest Mediator isolate discovered that exists in complex with liganded thyroid hormone receptor^7,12^, as well as of the positive cofactor 2 (PC2), a minimal form of the Mediator that mainly lacks the entire kinase module and many of the tail subunits while containing the metazoan-specific MED26. Interestingly, MED26 has a special requirement for Mediator-Pol II interaction in nuclear extracts (NE), which is still being investigated^6,13^.

Although these results implicated MED14 as a potentially critical subunit, the focus for Mediator-Pol II interactions in yeast was more on the Mediator head module subunits that were initially shown to be suppressors of otherwise lethal Pol II CTD truncation mutants^14,15^. In fact, these subunits have been implicated in interactions with Pol II based on cross-linking and structural studies. For instance, in the initial reports, while the Mediator middle module was observed to have contact points on Pol II’s RPB3 and RPB11 subunits^16^, the Mediator head module was also reported to interact with RPB4 and RPB7^2,17–21^, and MED17 with RPB3^22,23^. It was also shown that the RPB1 CTD might be responsible for the initial interactions as the deletion of RPB1 CTD 7-amino-acid sequence (heptad) repeats or insertion of a point mutation at every fourth residue of the CTD heptad repeats completely abolished Mediator interaction with Pol II^24^. This finding suggests that the RPB1 CTD may potentiate the Mediator-Pol II interaction, and this interaction could be further stabilized by the other Mediator and Pol II subunits mentioned above. Supporting the view that RPB1 represents Pol II’s primary interaction site with the Mediator, deletion of RPB4 and RPB7 subunits did not show a detectable decrease in Mediator-Pol II interaction^24^. Moreover, recombinant yeast Mediator head module, containing MED6, MED8, MED11, MED17, MED18, MED20, and MED22, was not sufficient to bind Pol II by itself and required other factors that included TFIIF^17,24,25^. In fact, the binding affinity of the head module to Pol II is very weak (almost 500-fold weaker as compared to full Mediator) based on surface plasmon resonance studies^24^, strongly suggesting that the critical interactions are yet to be identified. As the RPB1 CTD is not structured and is devoid of primary amines, cross-linking mass spectrometry, and cryo-EM approaches have also proven difficult to identify direct interacting partners^23,26^.

In our recent studies, using the MultiBac baculovirus expression system, we identified for the first time the composition of the minimal human core Mediator required for basal and p53-activated transcription. Our analyses of recombinant human Mediator subcomplexes showed that head (H), middle (M), and head + middle (H+M) modules could not interact with Pol II or support subsequent transcription, which was achieved only when MED14 was incorporated together with the head and middle modules (MED14+H+M)^6^. Our results pointing to a critical structural and functional role for MED14 were later verified by another group^26^.

In a significant extension of our previous study, we now report a primary recruitment mechanism for Pol II via the core Mediator complex, wherein the N-terminal half of MED14 (MED14-NTD) is critical for interaction with Pol II via the CTD of its RPB1 subunit and recruits it to core promoters to facilitate both basal and activated transcription. In addition, recombinant RPB1 but not ΔCTD RPB1 is sufficient to completely reverse the MED14-NTD+H+M-Pol II interaction. Recombinant MED14-NTD and full-length MED14 also interact directly with Pol II, although to a much lesser extent when compared with MED14-NTD+H+M. These results suggest that the initial Mediator-Pol II interactions are mediated through the MED14-NTD and the RPB1 CTD. Moreover, using the HHpred secondary structure prediction tool (https://toolkit.tuebingen.mpg.de/tools/hhpred), we show that the human MED14-NTD region (1-578 aa) has secondary structure features similar to yeast *S. pombe* MED14-NTD (1-548 aa)^27–31^ (https://zhanglab.ccmb.med.umich.edu/PSSpred/). Finally, a reanalysis of cryo-EM studies of *S. pombe* and *Saccharomyces cerevisiae* Mediator complexes also showed consistent results with our findings regarding the critical role of structurally conserved MED14-NTD in Pol II interaction through RPB1^23,26,32^.

## MATERIALS AND METHODS

### Reconstitution of Mediator subcomplexes and expression of Pol II subunits

Reconstitution of human Mediator subunits was performed as described previously^6^. Briefly, to co-express the human Mediator subunits, we used pFBDM transfer vectors into which the cDNAs encoding Mediator subunits were cloned^6^. Head (subunits: MED8, MED19, MED6, MED22, MED11, MED30, MED20, and MED18) and middle (subunits: HA:MED7, MED31, His:MED10, MED4, MED21, with or without MED26) modules were cloned into two separate pFBDM vectors. FLAG-tagged MED14-NTD (f:MED14-NTD), f:MED14-CTD, and f:MED14 (full MED14), as well as FLAG-tagged and non-tagged MED17 were also subcloned into separate pFBDM transfer vectors.

To produce the RPB subunits, total RNA from HeLa cells was purified, and cDNA was prepared by reverse transcription using oligo(dT) primers with Scientific Revert Aid First Strand cDNA Synthesis Kit (Thermo Fisher Scientific, Carlsbad, CA, USA; cat no: K1622), following the protocol recommended by the manufacturer. The resulting cDNA was amplified using appropriate PCR primers harboring His-or HA-tags to generate individual clones for each of the eleven RPB subunits (RPB2-RPB12). The f:RPB1, His:RPB1, and f:ΔCTD RPB1 constructs were prepared from pcDNA 3.1 (-) FLAG-Pol II WT plasmid (Addgene, Watertown, MA, USA; cat no: 35175) and cloned into separate pFBDM vectors.

To prepare baculoviruses, the pFBDM constructs were transformed into competent Dh10Bac cells for bacmid generation. The isolated bacmids were then used to transfect Sf9 cells for baculovirus production^33^. Expression of the proteins proceeded by infecting Hi5 cells at a density of 1 × 10^6^ cells/ml with the baculoviruses. Either single infection or co-infection was performed at this step, depending on the protein (or protein complex) to be produced. Cells were harvested at 60 hours post-infection by centrifugation at 1500 x g for 5 minutes. The pellet was resuspended in lysis buffer (500 mM KCl, 20 mM Tris-Cl [pH: 7.9], 20% glycerol [v/v], 0.1 mM EDTA [pH: 8.0], 3.5 mM β-mercaptoethanol, 0.1 mM PMSF, supplemented with the protease inhibitors pepstatin [0.5 μg/ml] and leupeptin [0.5 μg/ml]) and the cells were lysed mechanically using a Dounce homogenizer. The debris was removed by centrifugation at 13,000 x g for 15 minutes, and cell lysate was collected for further purification steps.

### Purification of proteins

All purified proteins outlined below were confirmed by western blot using chemiluminescence and either Coomassie or silver staining methods.

#### Purification of Mediator subcomplexes

Anti-FLAG M2 Affinity agarose beads (Sigma-Aldrich, St. Louis, MO, USA; cat no: A2220) were used to purify FLAG-tagged proteins. Beads were first washed with BC500 buffer (500 mM KCl, 40 mM HEPES-KOH [pH: 7.6], 0.4 mM EDTA, 0.5 mM PMSF, 0.5 mM DTT, and 0.1% NP-40), and the cell lysate was incubated with the beads overnight on a rotator at 4°C. Proteins were eluted with FLAG peptide (Sigma-Aldrich, St. Louis, MO, USA; cat no: F3290) and pooled. The eluate was applied to a gel filtration column packed with 10 ml Superose 6 (GE Healthcare, Uppsala, Sweden; cat no: GE17-0489-01) (ÄKTA purifier, GE Healthcare) and equilibrated with BC100 buffer (100 mM KCl, 40 mM HEPES-KOH [pH: 7.6], 0.4 mM EDTA, 0.5 mM PMSF, 0.5 mM DTT, and 0.1% NP-40). Finally, protein peak fractions corresponding to monomers-dimers (660 kDa) were pooled, frozen in liquid nitrogen, and stored at -80°C.

#### Purification of Pol II subunits (Rpb2-12)

Nickel beads were used for the purification of His-tagged proteins. Beads were first washed with BC300 buffer (300 mM KCl, 40 mM HEPES-KOH [pH: 7.6], 0.4 mM EDTA, 0.5 mM PMSF, 0.5 mM DTT, and 0.1% NP-40), placed and packed inside the column, and the lysate containing the protein of interest was loaded. The column was washed with BC250 buffer (250 mM KCl, 40 mM HEPES-KOH [pH: 7.6], 0.4 mM EDTA, 0.5 mM PMSF, 0.5 mM DTT, and 0.1% NP-40) containing 10 mM imidazole, and proteins were eluted with BC200 (200 mM KCl, 40 mM HEPES-KOH [pH: 7.6], 0.4 mM EDTA, 0.5 mM PMSF, 0.5 mM DTT, and 0.1% NP-40) containing 200 mM imidazole. HA-tagged proteins were purified with anti-HA agarose beads. The beads were first washed with BC300 and incubated with protein samples overnight. Following incubation, the beads were again washed with BC300, and the HA-tagged proteins were eluted with BC100 (100 mM KCl, 40 mM HEPES-KOH [pH: 7.6], 0.4 mM EDTA, 0.5 mM PMSF, 0.5 mM DTT) with HA peptide.

#### Purification of His:RPB1 and f:RPB1

The cells were harvested, and extracts were prepared as outlined above. A two-step purification process that includes size exclusion chromatography followed by affinity chromatography was used to purify f:RPB1 and His:RPB1. Briefly, the cell lysate (in BC300) was applied to a gel filtration column packed with 20 ml of Superpose 6 resin pre-equilibrated with BC300. RPB1-containing fractions corresponding to the monomeric/dimeric region were confirmed by western blot, pooled, and incubated with either M2 or nickel resin depending on the tag. The resulting eluates were re-confirmed by western blot.

#### Purification of f:ΔCTD RPB1

FLAG-tagged ΔCTD RPB1 was purified using Anti-FLAG M2 Affinity agarose beads. First, beads were washed with BC500 followed by overnight incubation with the cell lysate at 4°C, as described above. Proteins were eluted with BC100 buffer that contains FLAG peptide.

### Nuclear extract preparation from HeLa cells

Nuclear extracts were prepared using the method described by Dignam et al. with slight modifications^34^. Briefly, cells were harvested, washed with cold 1xPBS, and collected after centrifugation at 3000 x g for 5 minutes. Next, cells were suspended in two volumes of hypotonic buffer (10 mM HEPES [pH: 7.9], 1.5 mM MgCl_2_, 10 mM KCl, 0.5 mM DTT, 0.5 mM PMSF) and mechanically sheered using a Dounce homogenizer. Each 10-stroke set was followed by 10 minutes of incubation on ice (repeated three times). The homogenate was spun at 5000 x g for another 10 minutes. The nuclear pellet was re-suspended in equal volume of hypertonic buffer (20 mM HEPES [pH: 7.9], 25% glycerol [v/v], 0.42 M NaCl, 1.5 mM MgCl_2_, 0.2 mM EDTA, 0.5 mM PMSF, 0.5 mM DTT) and rocked at 4^°^C for 45 minutes. Finally, the homogenate was centrifuged for 15 minutes at 12000 g. The supernatant was collected as NE and was dialyzed to the desired salt concentration (BC100).

### Mediator immunodepletion

Mediator was immunodepleted from HeLa NEs as described previously^35^. Before immunodepletion, the anti-MED30 antibody was antigen purified as follows: *E. coli* BL21(DE3)pLysS bacteria transformed with pET11d-6His-MED30 plasmid were grown in two liters of LB broth until OD_600_ reached 0.4. Bacteria were then induced with 0.4 mM IPTG for four hours and harvested. The cells were suspended in denaturing lysis buffer (8 M ultrapure urea in 20 mM HEPES [pH: 7.9] and 500 mM NaCl), agitated slowly on a rotator at room temperature for 90 minutes, and centrifuged at 25,000 x g for 20 minutes. The pellet was discarded, and one bed volume of Ni-NTA-agarose (Qiagen Group; cat no: 30230) was added to the supernatant. The slurry was mixed and incubated for 90 minutes at room temperature. The slurry was packed into a column, and the resin was washed with 10 column volumes of denaturing lysis buffer. The bound protein was eluted with 100 mM EDTA in denaturing lysis buffer. Next, 0.5 g of CNBr-activated-Sepharose 4B (GE Healthcare, Uppsala, Sweden; cat no: GE17-0430-01) was swelled in 50 ml of 1 mM HCl. The matrix was equilibrated in denaturing lysis buffer and mixed with the purified recombinant protein in a 15 ml conical tube. After two hours of rotation at room temperature, the coupling reaction was blocked by adding Tris-HCl (pH: 8.0) to a final concentration of 0.1 M. Urea was gradually reduced to zero, and the trace amount of uncross-linked antigen was removed by washing the matrix with 5 ml of acidic wash buffer (0.1 M acetate [pH: 4], 0.5 M NaCl) and 5 ml of alkaline wash buffer (0.1 M Tris-HCl [pH: 8.0], 0.5 M NaCl), three rounds each. Finally, the column was equilibrated to a neutral pH with 10 mM Tris-HCl (pH: 7.4). Next, 10 ml anti-MED30 crude serum was diluted in 90 ml of 10 mM Tris-HCl (pH: 7.4) and applied to the MED30 cross-linked resin at a flow rate of 10 ml/h. After washing the column with 20 column volumes of 10 mM Tris-HCl (pH: 7.4), the antibody was eluted with 10 column volumes of 100 mM glycine (pH: 2.5) and 10 column volumes of 100 mM trimethylamine (pH: 11.5). The eluate was neutralized with one-tenth volume of 1M Tris-HCl (pH: 7.4).

To cross-link the Mediator antibody to protein A-Sepharose, 750 µl of protein A-Sepharose beads was first equilibrated with Tris-buffered saline. Three milligrams of the antigen-purified and pooled MED30-antibody from above were added and incubated with the beads overnight at 4^°^C. The next day beads were collected and washed with Tris-buffered saline followed by 0.2 M sodium borate (pH: 9.0). Beads were then resuspended in 7.5 ml of 0.2 M sodium borate (pH: 9.0), and 25 mg dimethyl pimelimidate (final concentration of 20 mM) was added. The suspension was rotated for 30 minutes at room temperature for cross-linking. The beads were washed with and incubated in 0.2 M ethanolamine for two hours at room temperature to stop cross-linking. Finally, the beads were washed with Tris-buffered saline and stored.

For immunodepletion of Mediator from HeLa NEs, the beads were equilibrated in BC200 buffer. Nuclear extract was adjusted to BC200 by adding KCl. The MED30-coupled protein A-Sepharose beads were mixed with the NE and incubated overnight at 4^°^C. Following incubation, the beads were centrifuged at 2000 x g for two minutes, and the Mediator depleted NE was collected for functional characterization studies.

### Mass spectrometry analysis

The samples were prepared by mixing the purified f:RPB1 with f:MED14+H+M, f:MED14-NTD+H+M, or f:MED14-CTD+H+M (500 nM each) and incubating the mixtures with anti-HA agarose beads for two hours (these complexes all contain HA-tagged MED7 subunit, as mentioned in the “Reconstitution of Mediator subcomplexes and expression of Pol II subunits” section above). The protein-coupled beads were washed with BC300 and sent to The Rockefeller University Proteomics Resource Center, where mass spectrometry analyses were performed.

Proteins were released from anti-HA agarose beads by on-bead trypsinization (Promega). The supernatant was reduced with DTT and alkylated with iodoacetamide, followed by second digestion. Samples were micro solid-phase extracted and analyzed by LC-MS/MS: 70-minute analytical gradient (2%B to 38%B), 12 cm built-in-emitter column, high resolution/high mass accuracy (Q-Exactive HF, Thermo). The data were processed using Proteome Discoverer 1.4 (Thermo Scientific) and searched using Mascot (Matrix Science) against UniProt human databases concatenated with common contaminants. Phosphorylation of serine, threonine, and tyrosine was allowed as a variable modification.

### *In vitro* transcription assays

*In vitro* transcription assays were performed as described previously^6,36^. Transcription reactions contained the following components: assay buffer (20 mM HEPES–KOH [pH: 8.2], 5 mM MgCl_2_, 60 mM KCl), 5 mM DTT, 0.5 mg/ml BSA, 20 units of RNasin, 1% polyethylene glycol, NTP mix (0.5 mM ATP, 0.5 mM UTP, 5 μM CTP, 0.1 mM 3′-O-methyl GTP), 10 µCi of [α-32P] CTP at 3000 Ci/mmol, 50 ng of DNA templates having G-less cassettes downstream of the adenovirus major late (ML) core promoter with enhancer sequences (5 x p53, 5 x Gal4, and 4 x ERE), and 50 µg of Mediator-depleted NE. Reactions were initiated by adding NE, Mediator subcomplexes (50 nM each), and activators (p53 and Gal4-VP16, 50 nM each) where indicated. Reactions were run at 30°C for 50 minutes, and the incorporation of radioactive CTP was squelched with 100 μM CTP for 20 minutes. Next, the reaction was terminated with the addition of stop mix (0.4 M sodium acetate [pH: 5.0], 13.3 mM EDTA, 0.33% SDS [v/v], 0.67 mg/ml yeast tRNA, 0.5 mg/ml Proteinase K). RNA products were isolated with phenol:chloroform:isoamyl alcohol (25:24:1) and precipitated in 100% ethanol, followed by 70%. Air-dried RNA was dissolved in formamide loading dye (95% formamide, 0.02% xylene cyanol, 0.02% bromophenol blue), resolved by electrophoresis on 5% polycrylamide-50% urea gels, and visualized via autoradiography. Band intensities were measured with ImageJ and used as an indicator of transcription strength^37^. Three independent repeats were performed for each transcription reaction.

### Immunoprecipitation

Immunoprecipitation analyses were performed as described previously^6^. Briefly, protein A-Sepharose beads were coupled with anti-MED6, anti-MED30, or anti-MED26 via incubating for three hours at 4°C. Beads were then washed with BC300, followed by BC150. Anti-MED6-, anti-MED30-, or anti-MED26-coupled protein A-Sepharose beads were incubated with recombinant Mediator variants (50 nM each) for three hours at 4°C. The protein-bound resin was washed with BC200 and further incubated with either purified RNA Polymerase II (50 nM) or individual RPB subunits (50 nM each) for three hours. The resin was washed with BC150. The immunoprecipitates were analyzed by immunoblotting using chemiluminescence.

### Immobilized template recruitment assays

The ML promoter region of the *in vitro* transcription assay template was amplified using 5’-biotinylated primers. The resulting biotinylated product was bound to Dynabeads M-280 Streptavidin (Invitrogen -Thermo Fisher Scientific, Carlsbad, CA, USA; cat no: 11205D). Briefly, the beads were washed with Binding & Washing (B&W) buffer (5 mM Tris-HCl [pH: 7.5], 1M NaCl, 0.5 mM EDTA), followed by incubation with the biotinylated templates in B&W buffer. Beads were again washed with B&W buffer containing 1 mg/ml BSA, %0.006 NP-40, followed by a further washing step with 1XPBS. Beads were then incubated in blocking solution (5 mM MgCl_2_, 20 mM HEPES-KOH [pH: 8,2], 5 mg/ml BSA, %0.03 NP-40, 12.5 mM DTT, 5 mg/ml PVP) for 15 minutes at room temperature followed by washing with wash buffer 1 (40 mM HEPES [pH: 7.5], 150 mM KCl, 4 mM MgCl_2_, 4 mM DTT, %0.1 NP-40). For each 150 µl reaction, 200 µg NE (mock- and Mediator-depleted extracts), the indicated Mediator subcomplex (200 nM), and 10 µg Poly (dI/dC) were mixed in assay buffer (20 mM HEPES–KOH [pH: 8.2], 5 mM MgCl_2_, 60 mM KCl), and incubated at 30°C for 50 minutes. Finally, beads were washed with wash buffer 2 (20 mM HEPES–KOH [pH: 8.2], 5 mM MgCl2, 100 mM KCl), and the bound sample was resolved with SDS-PAGE.

### Sources of antibodies and other proteins

Antibodies that were used in this study are as follows: MED14 antibody (Abcam, Cambridge, UK; cat no: ab170605), RPB1 N-20 (Santa Cruz Biotechnology, Dallas, TX, USA; cat no: sc-899), MED15 (Proteintech Group, Inc., Rosemont, IL, USA; cat no: 11566-1-AP), GST B-14 (Santa Cruz Biotechnology, Dallas, TX, USA; cat no: sc-138) and Anti-FLAG (Sigma-Aldrich, St. Louis, MO, USA; cat no: F7425).

Antibodies against other Mediator subunits and proteins (i.e., MED6, MED22, RPB1 8WG16, MED30, MED26, CCNC, MED28, MED13, MED12, p62, TAF100 [TAF5], MED7, MED4, Gdown1, RPB6, and RPB5) were obtained from Dr. Robert G. Roeder and were previously described and validated in western blotting experiments^6,13,38–41^. Anti-MED30 antibody that was affinity purified by chromatography against the bacterially expressed antigen^37^ was used for both Mediator immunodepletion of HeLa NEs and co-immunoprecipitation analyses.

GST:CTD and f:Gal4-VP16 proteins were obtained from Dr. Sohail Malik and Dr. Robert G. Roeder; f:p53 protein was obtained from Dr. Zhanyun Tang.

## RESULTS

### Reconstitution of WT and mutant MED14-core Mediator

Our previous results implicated the human core Mediator complex (MED14+H+M), but not the individual H or H+M modules, in the recruitment of Pol II to core promoters and the resulting transcription^6^. Here we aimed to dissect the underlying mechanism and test whether recombinant MED14 and its derivatives (NTD and CTD regions) could interact with Pol II independently, or the entire core Mediator assembly is necessary. Interestingly, the HHpred secondary structure prediction tool showed MED14-NTD as having an evolutionarily conserved secondary structure between human and *S. pombe* covering the KID, RM1, and RM2 regions (Figure 1a and Supplementary Table S1)^23,27^. This finding guided the design of the truncation mutations in Figure 1b for characterizing the conserved and non-conserved regions with our system.

**Figure 1.**
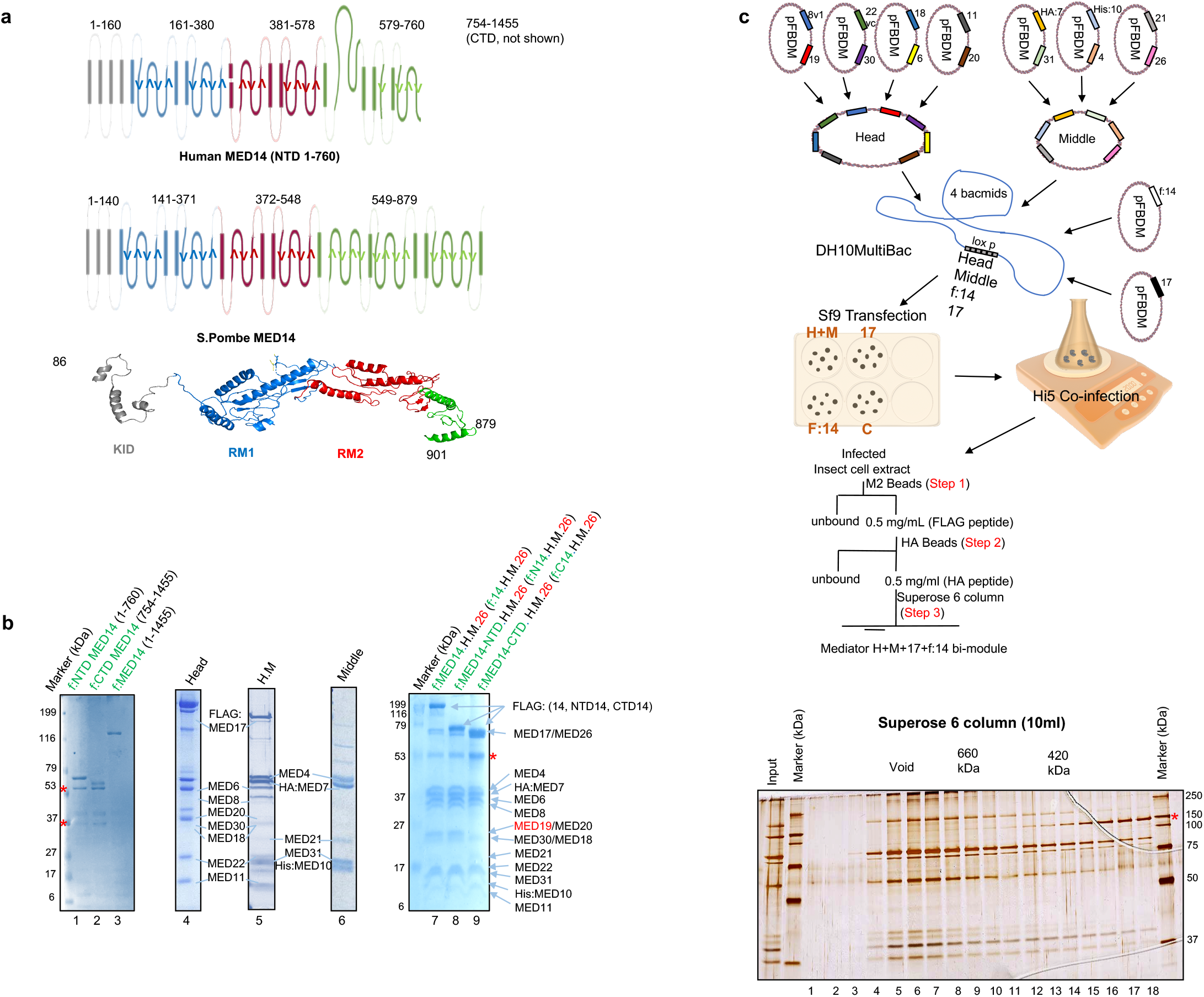
Purification of recombinant human MED14 deletion mutant fragments and reconstitution of Mediator subcomplexes. **(a)** HHpred secondary structure prediction and homology tool was used to identify evolutionarily conserved structured regions of yeast and human MED14 (also see Supplementary Table 1). The choice of the MED14 truncation site was based on these results. **(b)** SDS-PAGE (Coomassie blue staining) analysis of purified MED14 deletion mutants and Mediator subcomplexes. MED14 fragments in the amounts of 300 ng: f:MED14-NTD (1-760 aa), f:MED14-CTD (754-1455 aa), and f:MED14 (1-1455 aa) (lanes 1-3). Mediator subcomplexes in the amounts of 4 µg: H, H+M, M, MED14+H+M+MED26, MED14-NTD+H+M+MED26, and MED14-CTD+H+M+MED26 (lanes 4-9). All bands corresponding to the recombinant Mediator subunits are labeled. Red asterisks refer to non-specific proteins. MED19 is highlighted with red as its sub-stoichiometric. **(c)** A schematic for complex purification from (b) is shown. Mediator subcomplexes are expressed in insect cells and subsequently purified through M2 beads, HA beads, and Superose 6. Superose 6 column fractions were resolved in SDS-PAGE, and silver staining was performed. Red asterisks refer to non-specific proteins.

We therefore initially cloned, expressed (using baculovirus vectors in insect cells), and purified the FLAG-tagged MED14 derivatives that included full-length f:MED14 (1-1455 aa), the structurally conserved f:MED14-NTD (1-760 aa) region, and the non-conserved f:MED14-CTD (754-1455 aa) region (Figure 1b, lanes 1-3). Subsequently, Mediator subcomplexes containing the middle module (HA:MED7, MED31, His:MED10, MED4, MED21, with or without MED26), the head module (MED8, MED19, MED6, MED22, MED11, MED30, MED20, and MED18, as well as FLAG-tagged or non-tagged MED17), and the H+M modules were purified as described before (Figure 1b, lanes 4-6)^6^. Finally, 16-subunit human core Mediator complexes with MED14 derivatives were purified. These variants included H+M along with either f:MED14 (1-1455 aa), f:MED14-NTD (1-760 aa), or f:MED14-CTD (754-1455 aa) (Figure 1b, lanes 7-9). The purification steps, summarized in Figure 1c, yielded near stoichiometric amounts of all the constituent subunits, confirming complete assembly and homogeneity for each complex preparation. The gel filtration traces were followed on chromatography (Supplementary Figure S1). Interestingly MED19 composition in Med14-CTD+H+M+MED26 was lower compared to MED14+H+M+MED26 and MED14-NTD+H+M+MED26 (Figure 1b, lanes 7-9).

### MED14-NTD but not MED14-CTD-core Mediator is critical for RNA Pol II interaction

To test if MED14 and its mutant derivatives directly interact with Pol II, we first purified Pol II from HeLa NE. A subpopulation of Pol II in the metazoan cells contains a negative factor called Gdown1^42,43^. This form of Pol II (Pol II(G)) displays atypical Mediator requirements to overcome the negative effect exerted by Gdown1. To avoid any confounding results arising from this subpopulation, we ensured that we used a homogeneous preparation of standard Gdown1-free Pol II. Immunoblot analyses confirmed that following purification over different chromatographic steps, our Pol II was mostly devoid of Gdown1 (Supplementary Figure S2a). Next, we performed co-immunoprecipitation assays with the 8WG16 anti-Pol II RPB1 CTD antibody. Interestingly, MED14-NTD and full-length MED14, but not MED14-CTD, interacted with Pol II (Supplementary Figure S2b), indicating that MED14, through the NTD, can intrinsically interact with Pol II. Since MED14 and MED14-NTD alone showed only relatively weak interactions with Pol II, we repeated the co-immunoprecipitation reactions with MED14, MED14-NTD and MED14-CTD assembled into core Mediator complexes along with H and H+M modules to see if the complete assembly of core Mediator would increase the interaction with Pol II. Antibody against a core subunit, MED6, was used for the co-immunoprecipitation reactions. Due to the conditional requirement of MED26 for core Mediator-Pol II interaction in NE, we also included MED26 in our constructs when necessary^6,13^. We did not observe any interaction of Pol II with the purified H and H+M (Figure 2a, lanes 2-3), consistent with our earlier report^6^. In contrast, with the inclusion of MED14, we detected strong Pol II interactions exclusively with MED14+H+M+MED26 and MED14-NTD+H+M+MED26 complexes, but none with the MED14-CTD+H+M+MED26 complex (Figure 2a, lanes 4-6), demonstrating a critical role for the structurally conserved N-terminal region of MED14 in the Pol II interaction. In addition, MED14-NTD+H showed weaker interaction with Pol II compared to MED14-NTD+H+M+MED26, again suggesting the necessity of the complete core structure for maximum Pol II interaction (Figure 2b, lane 4 vs. lane 5). In this NE-free pull-down system, MED26 demonstrated no effect on core Mediator-Pol II binding (data not shown).

**Figure 2.**
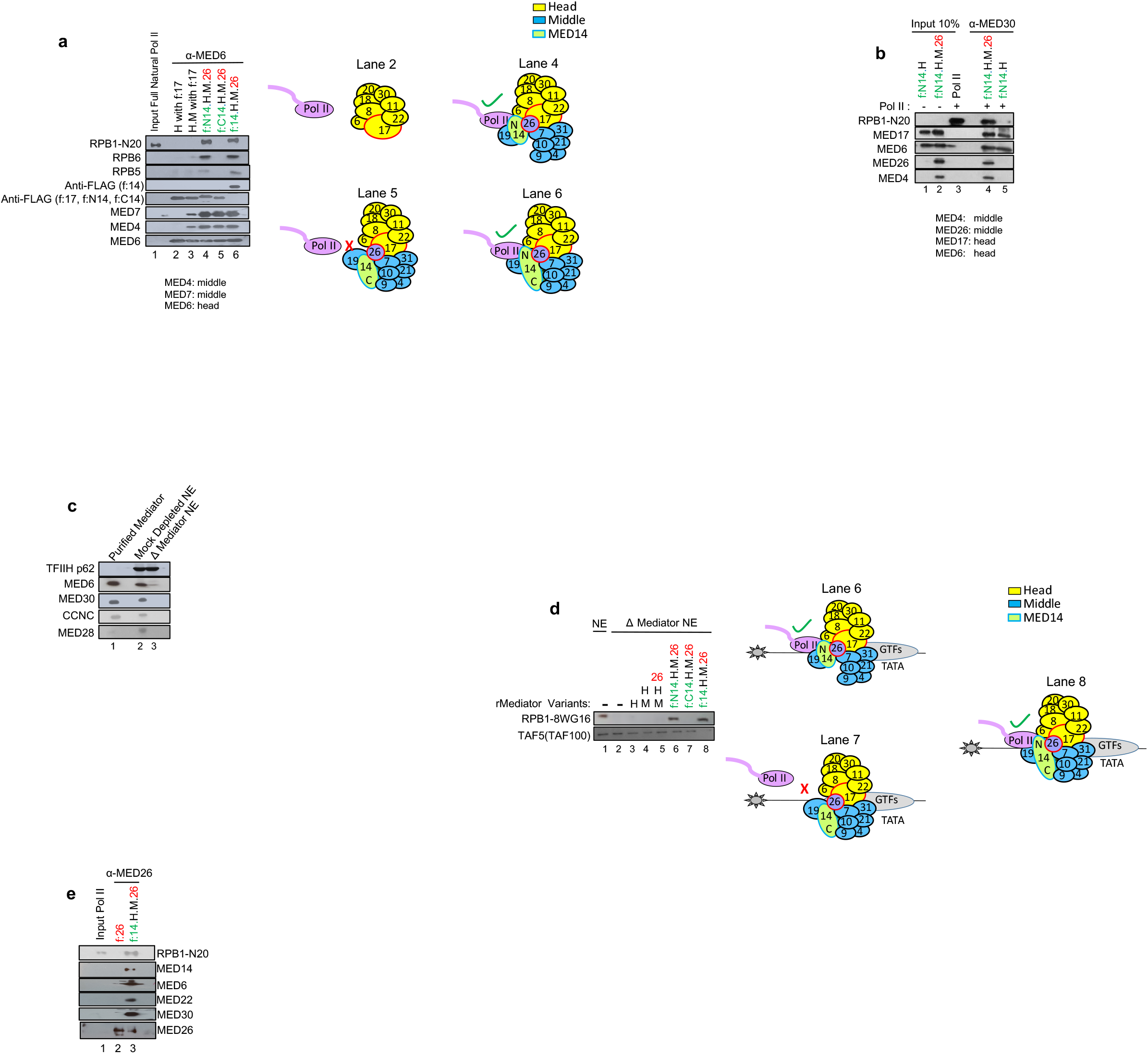
MED14-NTD-containing human core Mediator interacts with RNA Polymerase II and recruits it to AdML promoters. **(a)** Western blot analysis of recombinant Mediator and Pol II interaction assay. Equimolar concentrations (50 nM) of either one of the recombinant human Mediator H, H+M, MED14-NTD+H+M+MED26, MED14+H+M+MED26, or MED14-CTD+H+M+MED26 subcomplexes were independently conjugated to α-MED6-coupled beads. Subsequently, 50 nM of Pol II was added to each reaction, and the co-purified proteins were assayed by western blot. Lane 1 shows the input of purified Pol II, and lanes 2-6 show the precipitated proteins indicating interactions with the bead-conjugated Mediator subunits. Two anti-FLAG blots are shown. The upper blot represents f:MED14 (170kDa), while the lower one shows f:MED17, f:MED14-NTD, and f:MED14-CTD (all ∼75 kDa). Diagrams of Mediator subunit composition for H, H+M, MED14-CTD+H+M+MED26, MED14-NTD+H+M+MED26, and MED14+H+M+MED26 and their interaction profiles with Pol II are also shown. **(b)** MED14-NTD+H+M+MED26 interacts with Pol II. Immunoprecipitation was repeated as in (a) with α-MED30 antibody. **(c)** Mediator was immunodepleted from NE using anti-MED30 antibody. Western blot of purified natural Mediator, mock-depleted NE, and Mediator-depleted NE is shown. **(d)** Immobilized template assay for Pol II recruitment by recombinant Mediator subcomplexes is shown. Biotinylated DNA template containing a core promoter was incubated with either control (none-depleted HeLa NE) (lane 1) or Mediator-depleted HeLa NE (lanes 2-8) together with recombinant Mediator variants (200 nM each). Recruitment of Pol II (RPB1) and TFIID (TAF100) was monitored by western blot. Representative diagrams for the Pol II recruitment are shown. **(e)** MED26 does not directly interact with Pol II. The experiment in (a) was repeated using either f:MED26 alone or f:MED14+H+M+MED26 incubated with Pol II. Immunoprecipitation was done using α-MED26.

Next, to test whether the observed increase in Mediator-Pol II interactions also leads to increased recruitment of Pol II to core promoters, we characterized the recruitment of Pol II to ML promoter by the human core Mediator subcomplexes using immobilized template assays with biotinylated DNA template beads. In this assay, we used a competing DNA analog (dI/dC) to minimize the non-specific binding of proteins. We incubated bead-immobilized DNA with HeLa NE that contained all PIC components, including GTFs and Pol II, but (almost) no Mediator, which had been specifically depleted using an antibody against MED30 (Figure 2c). Pol II in Mediator-depleted NE is known to show defects in recruitment to core promoters^6,38^. We, therefore, incubated Mediator-depleted NE with the recombinant Mediator subcomplexes from Figure 1b to assess the recovery of Pol II recruitment to promoters. For this purpose, PIC formation was monitored by immunoblotting of Pol II and selected GTFs. As expected, TFIID (monitored by TAF5), which has a general intrinsic affinity for DNA, was bound to DNA independent of the Mediator. Most importantly, none of the H, H+M, H+M+MED26, or MED14-CTD+H+M+MED26 complexes that failed to interact with Pol II recruited Pol II to promoters (Figure 2d, lanes 3-5, and 7). In contrast, the MED14+H+M+MED26 and MED14-NTD+H+M+MED26 complexes fully recruited Pol II to the core promoter at levels comparable to endogenous Mediator (Figure 2d, lanes 1, 6, and 8). As we observed a MED26 requirement for MED14-NTD+H+M-Pol II interaction in NE, but not in the purified system, we further tested whether MED26 alone could interact directly with Pol II. We again saw the dramatic interaction of MED14+H+M+MED26, but not MED26 alone, with Pol II (Figure 2e). The interaction was dependent on MED14+H+M (Figure 2e)^6^. The specific requirement of MED26 for Pol II interaction in NE is currently being investigated.

### RPB1 interacts with MED14-NTD+H+M through its CTD domain

To determine the Pol II subunit(s) required for the MED14-NTD+H+M+MED26 interaction, they were individually expressed as recombinant proteins (Supplementary Figure S3a-d) and analyzed in co-immunoprecipitation assays. We observed significant interaction of RPB1, but not other isolated RPB subunits, with MED14-NTD+H+M (Figure 3a-d and Supplementary Figure S4a-f). Interestingly, mass spectrometry results done with MED14-NTD+H+M-RPB1 complex revealed the hypo-phosphorylated version of RPB1 to interact with MED14-NTD+H+M (Figure 3a, right panel). Having narrowed down the interaction to RPB1, we further dissected its CTD and non-CTD (ΔCTD RPB1) domains. Co-immunoprecipitations with antibodies against the MED30 subunit of the reconstituted Mediator subcomplexes showed direct interactions of RPB1 and GST:CTD, but not ΔCTD RPB1, with MED14-NTD+H+M (Figure 3b, left and right panels). Moreover, MED14+H+M and MED14-NTD+H+M interaction with RPB1 gave a strongly associated 16-subunit complex (with or without MED26) (Figure 3a, Figure 3e, and Supplementary Figure S4a). We next tested if recombinant RPB1 and intact Pol II would compete for binding to MED14+H+M+MED26. Interestingly, recombinant His:RPB1 could reverse the MED14-NTD+H+M+MED26-Pol II interaction by at least 50% when supplemented in a 1:1 ratio (Figure 3f, left panel). More importantly, when supplemented in excess, recombinant RPB1 but not ΔCTD RPB1 could completely reverse the MED14-NTD+H+M+MED26-Pol II interaction (Figure 3f, right panel), once again suggesting that RPB1, acting through its CTD, might be the primary subunit that binds to MED14-NTD+H+M+MED26 and that these subunits potentiate further interactions between H+M and Pol II.

**Figure 3.**
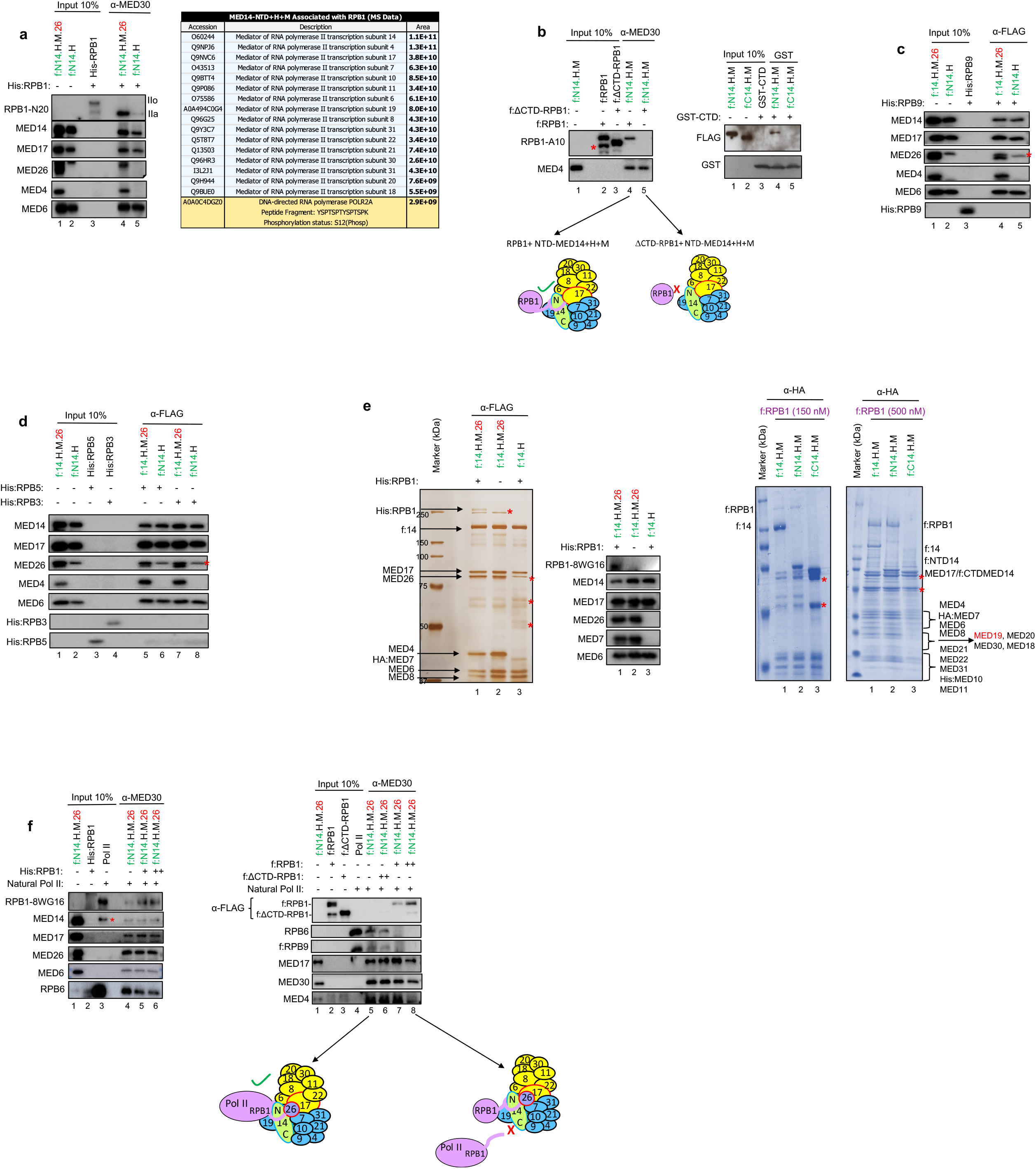
RPB1 CTD interacts directly with MED14-NTD-core Mediator. **(a)** *In vitro* interaction assay for MED14-NTD+H+M+MED26 and RPB1 is shown. We used 50 nM of proteins or protein complexes for all binding assays. MED14-NTD+H+M+MED26 and MED14-NTD+H were conjugated to beads through α-MED30. Recombinant His:RPB1 (50 nM) was added to each reaction, and the co-purified proteins were assayed by western blot. Only the hypo-phosphorylated (IIa) form of RPB1 was detected, as also confirmed by mass spectrometry (right panel). **(b)** Interaction assay was repeated as in (a) with f:RPB1, f:ΔCTD RPB1 (left panel), and GST:CTD (right panel). Red asterisk shows degraded RPB1 protein. The interaction is also represented in the diagram. **(c-d)** Binding reactions were repeated as in (a) with recombinant His:RPB9, His:RPB5, and His:RPB3 subunits. Red asterisk shows MED17 antibody from prior blotting that could not be completely stripped off. **(e)** Silver stain image (leftmost panel) shows co-purification of His:RPB1 and f:MED14+H+M+MED26 (lane 1). Lysates of insect cells separately expressing His:RPB1 and those expressing either f:MED14+H+M+MED26 (lane 2) or f:MED14+H (lane 3) were mixed and purified with M2 agarose beads targeting f:MED14. The resulting eluates were run on SDS-PAGE, silver stained, and verified via western blot (shown next to the silver staining). Interaction of f:RPB1 (150 nM and 500 nM) with f:MED14+H+M, f:MED14-NTD+H+M, and f:MED14-CTD+H+M (500 nM each) was also checked using anti-HA agarose beads (mass spectrometry-verified Coomassie stained gels, rightmost two panels). Red asterisks show non-specific protein bands. **(f)** RPB1 competition assay with Pol II for binding to MED14-NTD+H+M+MED26 is shown. The experiment in (a) was repeated with MED14-NTD+H+M+MED26 in the presence of equimolar amounts of RPB1 as competitor. Either equal amounts of Pol II and His:RPB1 (left panel) or up to 10-fold more f:RPB1 and f:ΔCTD RPB1 relative to the level of Pol II (right panel) were added to the reaction. A schematic representation of this competition assay is illustrated next to the right panel. Red asterisk shows a non-specific protein band.

### MED14-NTD is required for basal and activated transcription

To extend our mechanistic studies to function, we next performed *in vitro* transcription assays. In this assay, we used the Mediator-depleted NE from Figure 2c. This extract does not yield specific RNA products and thus is transcription-defective unless the Mediator is added back. Therefore, we checked if the addition of recombinant Mediator subcomplexes was sufficient to stimulate transcription in these Mediator-depleted NEs. We first assessed recovery of basal transcription with the recombinant human core Mediator subcomplexes described in Figure 1b. Two conventional core promoter-containing DNA templates that yield different-size RNA products (AdML1 and AdML2) were used per reaction. As expected from mechanistic studies performed in Figure 2 and Figure 3, and remarkably, MED14-NTD+H+M+MED26 fully recovered basal transcription (Figure 4a, lane 7 vs. lanes 3-6 and 8). In contrast, H, M, H+M, H+M+MED26, and MED14-CTD+H+M+MED26 subcomplexes showed transcription levels comparable to the negative control, indicating that they have no effect on the recovery of transcription (Figure 4a, lane 2 vs. lanes 3-6 and 8). Mediator is known to be recruited by many activators through interactions with its tail module, the p53-mediated recruitment through a p53-MED17 interaction being a notable exception^7,44^. This afforded us an opportunity to also extend our studies to assess whether MED14-NTD+H+M+MED26 could support activator-dependent transcription. For this purpose, we repeated our in vitro transcription assays with purified p53 and, as a control, with MED25 (tail)-interacting Gal4-VP16 (Figure 4b-c)^1,45,46^. Recombinant MED14-NTD+H+M+MED26, as well as the endogenous intact Mediator complex, fully responded to the p53 activator (Figure 4c, lanes 1-2, lanes 7-8 and 13-14). This finding is consistent with the notion that p53 recruits Mediator through MED17^7,44^. We repeated the same experiment with another activator, Gal4-VP16. In this case, the fold-differences between basal and activated transcription mediated by MED14+H+M+MED26 (Figure 4d, lanes 17-18) and MED14-NTD+H+M+MED26 (Figure 4d, lanes 13-14) are relatively weaker compared to that observed with the natural Mediator (Figure 4d, lanes 1-2 and 19-20). This result is consistent with the notion that Gal4-VP16 requires a tail module subunit for maximal transcription, which is absent in our reconstituted core complex^1,45^. Gal4-VP16 supplemented HeLa NE also gave a weak background transcript level, most probably due to the residual Mediator complex that was still present in the Mediator depleted HeLa NE (Figure 4d, lanes 4, 6, 8, 10, 12, and 16). Overall, MED14-NTD+H+M (with or without MED26) not only interacts with Pol II via the RPB1 CTD (Figure 2 and Figure 3) but also mediates functional Pol II recruitment to core promoters (Figure 2d) that in turn facilitates basal and activator-driven transcription (Figure 4a, 4c, and 4d).

**Figure 4.**
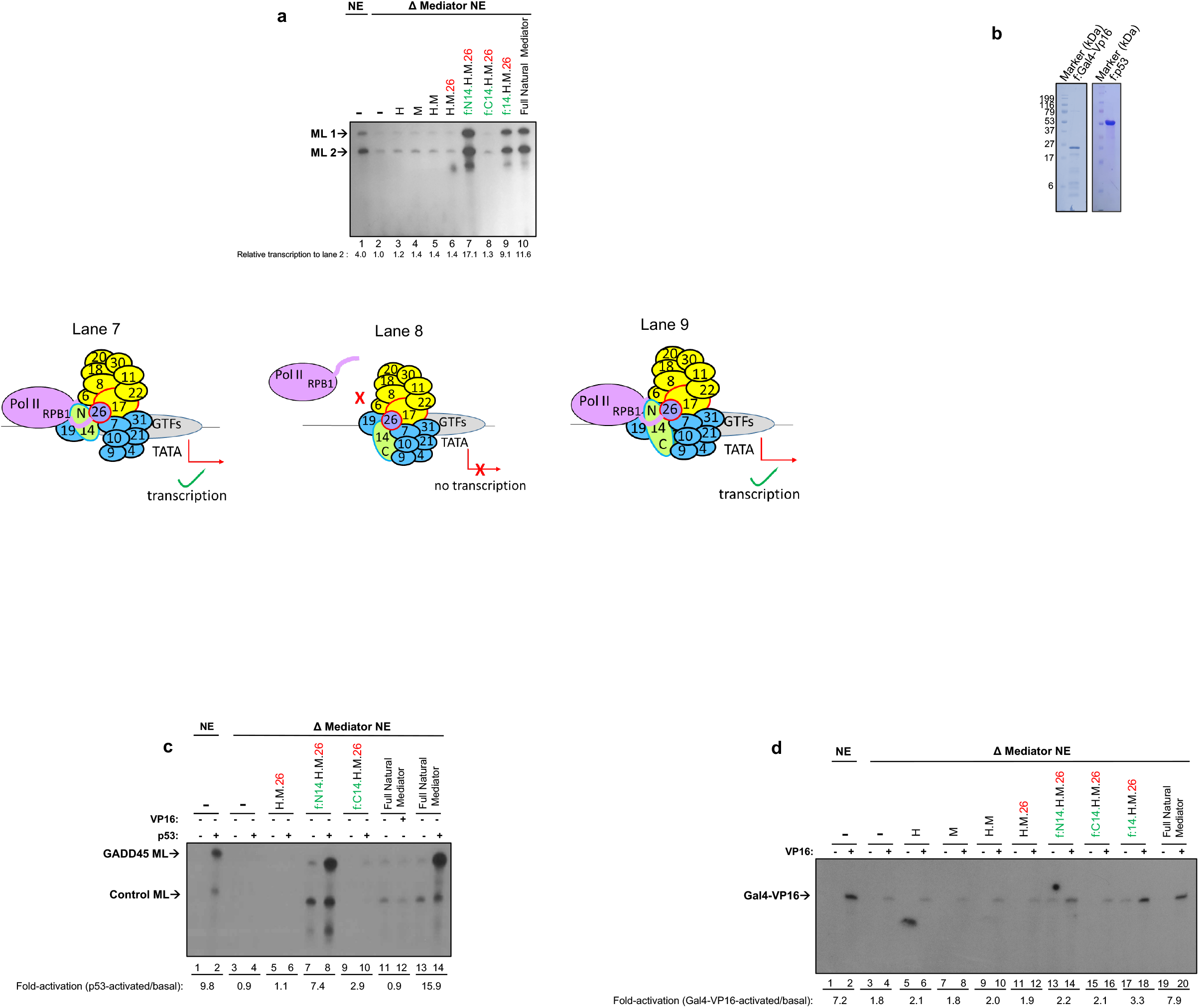
MED14-NTD is necessary for a full recovery of basal and activated (p53) transcription. **(a)** *In vitr*o transcription with Adenovirus ML-containing core promoters was performed. The reactions were carried out with Mediator-immunodepleted HeLa NE supplemented with recombinant Mediator complexes from Figure 1b as indicated. RNA products are annotated as ML1 and ML2. The band intensities of the RNA products are measured, the mean intensities of the two RNA product bands on each lane - ML1 and ML2 – are calculated, and the values are normalized to lane 2 (Mediator-immunodepleted HeLa NE without recombinant Mediator complex). The experiment is repeated three times and a representative gel is depicted (the quantitation under the gel is for the particular experiment shown). A schematic representation of the *in vitro* transcription assay is also illustrated. **(b)** Purified recombinant Gal4-VP16 and p53 proteins (Coomassie stain). **(c)** Autoradiogram of *in vitro* transcription as in (a) using p53 and its cognate template GADD45. RNA products are labeled as GADD45 ML (p53 responsive) and control ML (not responsive to p53 -basal transcription-). The band intensities of the RNA products (upper bands, where GADD45 was used) were quantified using ImageJ, and data were expressed as fold-changes (activated vs. basal transcription) for each Mediator complex derivative. Lane 12 contained Gal4-VP16 to serve as a negative control. The experiment is repeated three times and a representative gel is depicted (the quantitation under the gel is for the particular experiment shown). **(d)** Autoradiogram of *in vitro* transcription as in (c) supplemented with Gal4-VP16 and its cognate template. RNA products are labeled as Gal4-VP16. The experiment is repeated three times and a representative gel is depicted (the quantitation under the gel is for the particular experiment shown).

## DISCUSSION

The large multi-subunit Mediator serves as a critical coactivator for functional communication between enhancer-bound transcriptional activators and the promoter-associated general transcription machinery. Toward an understanding of underlying mechanisms and the functions of individual Mediator subunits and modules, we previously employed the MultiBac system to generate a recombinant core Mediator capable of carrying out the primary effector function of the Mediator and further identified a pivotal role for MED14 subunit in the associated Mediator-Pol II interactions. This approach opened new possibilities towards characterizing the Mediator not only at the modular level but also at subunit and domain levels. Importantly, this functional core showed all the fundamental hallmarks of the natural Mediator complex related to Pol II interaction and function^6^. Thus, in a significant extension of our previous study and using the same approach, here we shed light on the critical roles of Mediator in the formation of a functional PIC. By recombinantly generating Mediator subcomplexes and truncation mutants of MED14 and RPB1, we describe a mechanism critical for physical and functional interactions of the human core Mediator complex (MED14+H+M) with Pol II. Here, through multiple lines of evidence, we report the essentiality of the structurally conserved NTD region of MED14 (1-578 aa, covering KID, RM1, and RM2 domains) for Pol II interaction via the RPB1 CTD and for subsequent transcription (see the model in Figure 5). Our results are consistent with previous studies in yeast that highlight the MED14-NTD as a critical region for basal transcription^9–11^.

**Figure 5.**
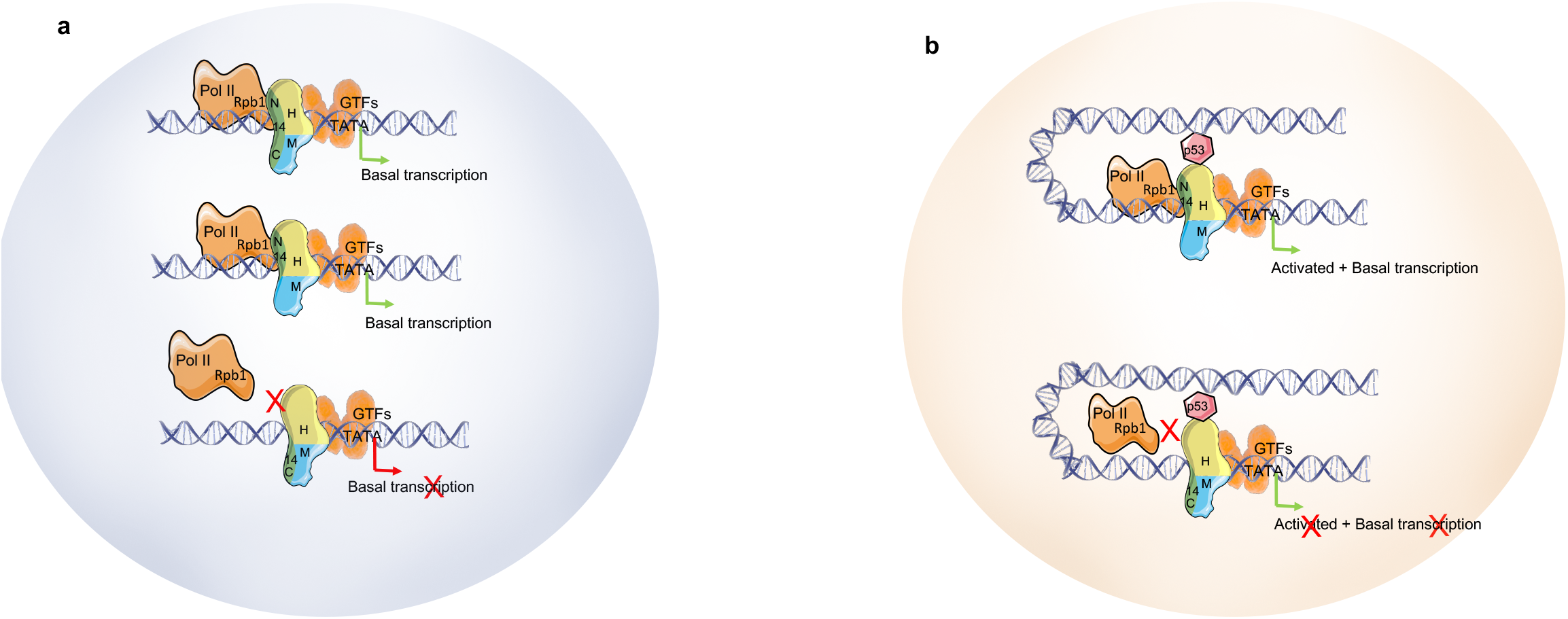
Summary: MED14-NTD-containing core Mediator interaction with RPB1 of Pol II facilitates transcription *in vitro*. **(a-b)** Pol II interacts through its RPB1 CTD with MED14-NTD+H+M *in vitro*. This MED14-NTD-containing core, in turn, recruits Pol II to core promoters to facilitate both basal **(a)** and activated **(b)** transcription (p53). The model is presented on nucleosome-evicted (accessible) DNA.

We also revisited yeast Mediator-Pol II cryo-EM structures previously deposited by others in the PDB database (https://www.rcsb.org/) to check if there was any evidence for MED14-NTD and RPB1 CTD interaction^23,26,32^. Unfortunately, most of the CTD repeats of the RPB1 region were omitted in the structural studies, most likely because they were disordered. Moreover, MED14 characterization was also omitted in other structural studies^24^. Nonetheless, our reanalysis revealed clear interactions between MED14-NTD and residual RPB1 CTD in the structure (Supplementary Figure S5, PDB 5U0S)^47^. In fact, and importantly, the MED14-NTD region in the 15-subunit core Mediator showed proximity (within a few Angstroms) to the residual CTD domain of Pol II, again suggesting that the initial interaction could be facilitated by these regions (Supplementary Figure S5)^23,26,32^. As most of the RPB1 CTD was not mapped in these cryo-EM studies, a possible strong interaction extending beyond a few amino acids was not observed (PDB IDs: 5U0S, 5N9J, and 5SVA)^23,24,26,32,47,48^.

What might be the underlying mechanistic basis for this functional requirement of MED14-NTD interaction with RPB1 CTD? It was previously suggested that involvement of the RPB1 CTD in holoenzyme formation might be a prelude to a more extensive set of interactions that occur later in the PIC formation pathway^1–4,8,9^. However, the persistent role of MED14-NTD in RPB1 CTD interaction has not been ruled out, and this may have important functional consequences (see below).

In the previous structural studies, head and middle modules of the Mediator have been reported to interact with RPB3, RPB4, RPB7, and RPB11 subunits of Pol II^2,16–23^. Moreover, deletion of RPB1 CTD completely abolished interaction of Mediator with Pol II, raising the possibility that the initial recruitment of Pol II is through its CTD domain^24^. Interestingly, although we report here that MED14-NTD is necessary for the overall interaction of Mediator with Pol II, this interaction is maximal when MED14-NTD is incorporated into the H+M module. Moreover, MED14-NTD+H+M interaction with RPB1 is abolished when RPB1 CTD is deleted. This finding suggests that the herein-described MED14-RPB1 contacts may be the initial interaction points between Mediator and Pol II and that they potentiate further interactions between H+M and Pol II. Alternatively, it is also possible that upon binding to H+M, MED14-NTD assumes a proper conformation for a maximal Pol II interaction and function.

The RPB1 CTD is known to be differentially phosphorylated, especially at Ser2 and Ser5, as transcription progresses from initiation to elongation^49,50^. It is tempting to speculate that CTD phosphorylation also depends on MED14 interactions as this subunit is critical for the recruitment of Pol II to promoters. It is equally possible that MED14-RPB1 CTD interaction is dynamic and depends on the phosphorylation state of CTD.

Our present results clearly show that MED14-CTD and MED14-NTD could interact with H+M independently. Moreover, our previous cross-linking mass spectrometry results revealed contacts indicative of potential looping between the MED14-CTD and MED14-NTD regions^6^. As the MED14-CTD is bound to tail module subunits, it is plausible that this looping may hinder MED14-Pol II binding through the intervention of tail subunits in the absence of activators, thus affecting the subsequent transcription. Once enhancer-bound activators bind to tail subunits, the conformational changes could facilitate MED14-Pol II interactions leading to transcription. This possibility will be tested in the future using our reconstitution approaches and biochemical assays as we start to assemble the tail module subunits onto our existing human core Mediator.

Our reconstitution approaches also suggest a possible differential regulation model for various activators that bind to the Mediator complex. For instance, the tail module subunits that assemble with the Mediator through the MED14-CTD are the primary targets for activators. On the other hand, certain activators, such as p53, are known to bind to other components of the Mediator, including core subunits^44^. While the looping of MED14-CTD onto MED14-NTD may regulate typical activators through tail module intervention for Mediator-Pol II interaction, a subset of activators, like p53, that target core Mediator subunits may not be affected by this mechanism and thus could be regulated differently. This opens the possibility of designing selective modulators of the p53-core Mediator interface without affecting other tail-activator interactions. Importantly, these hypotheses are readily testable using our reconstitution approaches and biochemical techniques.

Finally, structural studies, including cryo-EM and cross-linking mass spectrometry analyses, have been seminal in characterizing the interaction between Pol II and Mediator. However, studying Mediator-Pol II functional interactions has remained challenging due to the large size of these complexes, lack of primary amines at critical sites (which renders such sites invisible in cross-linking mass spectrometry), and the dynamic nature of the interactions. As a result, the critical role of MED14 in Pol II interactions and subsequent transcription is overlooked. We believe our biochemical reconstitution analyses will provide a vital addition to the tools for studying Mediator functions.

## AVAILABILITY

The secondary structures of human and *S. pombe* MED14-NTD regions were compared using HHpred. HHpred is an online tool for protein structure prediction and homology analysis, which can be accessed at: https://toolkit.tuebingen.mpg.de/tools/hhpred

The protein structure/interaction maps referred to in the text are available online on The RCSB Protein Data Bank (https://www.rcsb.org/) with the PDB IDs: 5U0S, 5N9J, and 5SVA.

## ACKNOWLEDGEMENT

We are grateful to Dr. Robert G. Roeder and Dr. Sohail Malik for their critical inputs, discussions, and antibodies. We thank Dr. Keiichi Ito, Dr. Mesut Muyan, and Dr. Takashi Onikubo for their critical inputs to the writing of the manuscript. We thank Dr. Timothy Richmond for the MultiBac baculovirus system. We thank the Library of Sciences and Medical Illustrations and Smart PPT for illustrative figures. The Rockefeller University Proteomics Resource Center acknowledges funding from The Leona M. and Harry B. Helmsley Charitable Trust and Sohn Conferences Foundation for mass spectrometer instrumentation.

## Supplementary Figure Legends

**Supplementary Figure S1. Gel filtration traces for recombinant core Mediator subcomplexes**

Recombinant complexes are purified as depicted in Figure 1b-c and run over 10 ml of Superose 6 gel filtration column. Chromatography traces of three core Mediator subcomplexes are represented.

**Supplementary Figure S2. Purification of Gdown1-free Pol II and MED14 interaction with Pol II**

**(a)** Purification of Gdown1-free Pol II. Nuclear extract from HeLa cells stably expressing f:RPB9 was run on HiTrap Q HP (anion-exchange, step 1) followed by HiTrap Heparin (cation-exchange, step 2) columns. Pol II fractions mostly cleared from Gdown1 were pooled together and further purified over M2 agarose affinity resin (step 3). The right panel shows SDS-PAGE followed by silver staining of purified Pol II. **(b)** Western blot analysis of recombinant MED14 and Pol II interaction assay. Equimolar amounts of Pol II (50 nM) were conjugated to α-8WG16-coupled (against RPB1) beads. 50 nM each of recombinant f:MED14-NTD, f:MED14-CTD, and f:MED14 was added to each reaction, and the co-purified proteins were assayed by western blot. Lanes 1-3 show the input of recombinant MED14 fragments. Lane 4 shows the input of Pol II. Lanes 5-7 show the precipitated proteins indicating interactions with the bead-conjugated Pol II.

**Supplementary Figure S3. Expression of FLAG-, His-, and HA-tagged RPB subunits in insect cells**

**(a)** SDS-PAGE (Coomassie staining) analysis of purified f:RPB1. **(b)** Superose 6 gel-filtration spectra and the corresponding western blot of His:RPB1 eluates. **(c)** Monomer and dimer eluates of His:RPB1 from (b) were pooled together and further purified using Nickel resin. **(d)** Western blot of individually cloned and purified His:RPB3, His:RPB5, His:RPB6, His:RPB7, His:RPB9, His:RPB12, HA:RPB4, HA:RPB8, HA:RPB10, and HA:RPB11.

**Supplementary Figure S4. Characterization of recombinant Pol II subunit interactions with reconstituted human core Mediator subcomplexes**

**(a-f)** Western blot analysis of recombinant Mediator subcomplexes and RNA Pol II subunit interaction assays. Equimolar amounts (50 nM each) of either recombinant human Mediator, MED14-NTD+H+M+MED26, MED14+H+M+MED26, or MED14-NTD+H were conjugated to either α-MED30-**(a)** or α-FLAG-coupled **(b-f)** beads. Recombinant RPB subunits from Supplementary Figure S3 were added to each reaction, and the co-purified proteins were assayed by western blot. Red asterisks show MED17 antibody from prior blotting that could not be completely stripped off.

**Supplementary Figure S5. Pol II-MED14 interaction as illustrated on PDB Database**

PDB ID: 5U0S revealed the MED14-NTD region (RM1) to be in close proximity with the CTD linker region of RPB1. Most of the disordered CTD was not characterized in the structure.

**Supplementary Table S1. HHpred secondary structure prediction of *S. pombe* and human MED14**

Sequence and secondary structure similarities between *S. pombe* and human MED14 were assessed using HHpred.

