## Supplementary Figures 1-5 for "Mediator-RNA Polymerase II Interactions Critical for Transcriptional Activation Are Mediated by the N-terminal Half of MED14 and C-terminal Domain of RPB1"

Gel Filtration Traces of f:MED14+H+M+MED26, f:MED14-NTD+H+M+MED26 and f:MED14-CTD+H+M+MED26

**Supplementary Figure S1**

**Superose 6 column (10ml)**

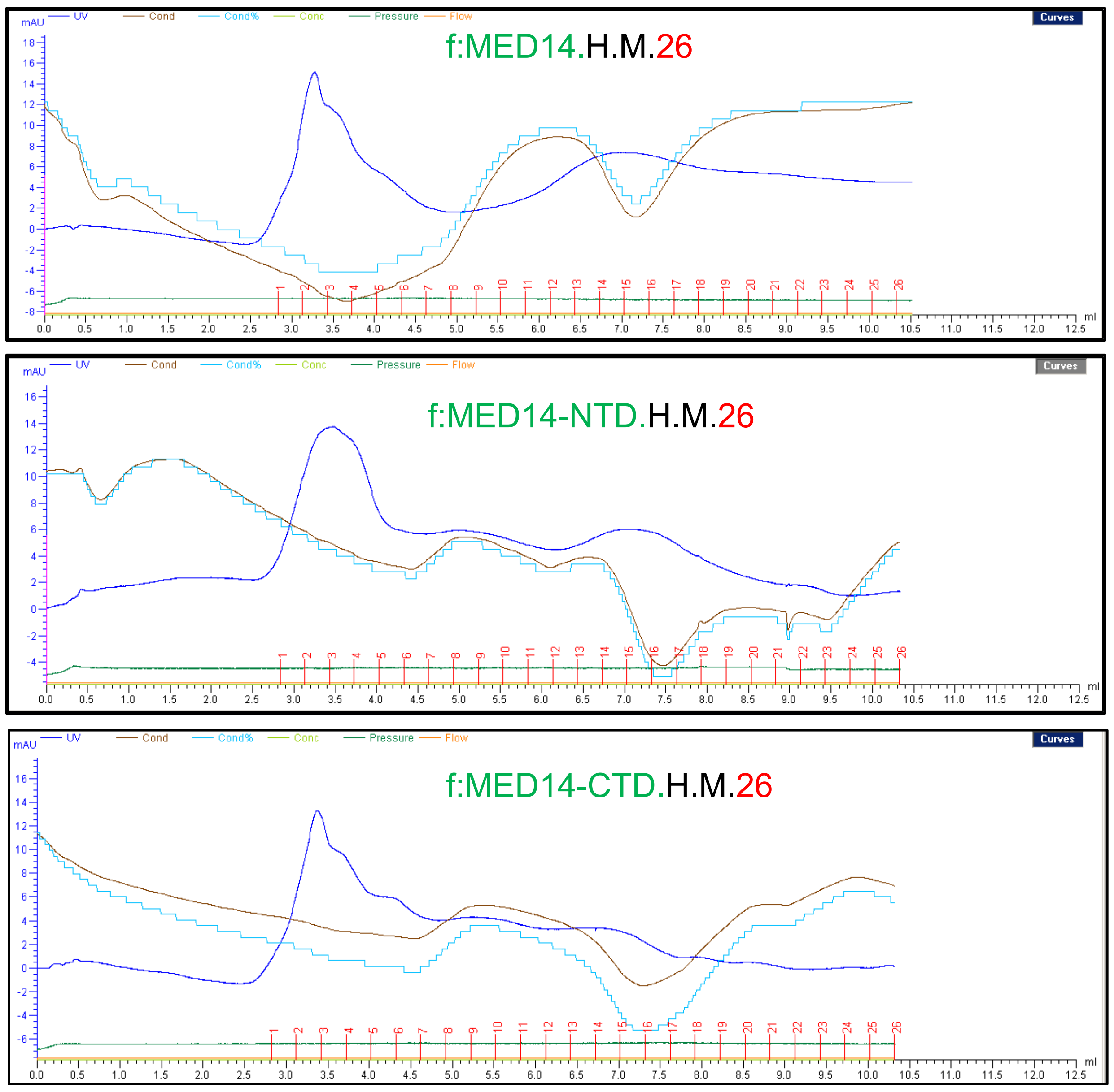

Purification of Pol II and IP with Recombinant MED14 Variants

### Supplementary Figure S2

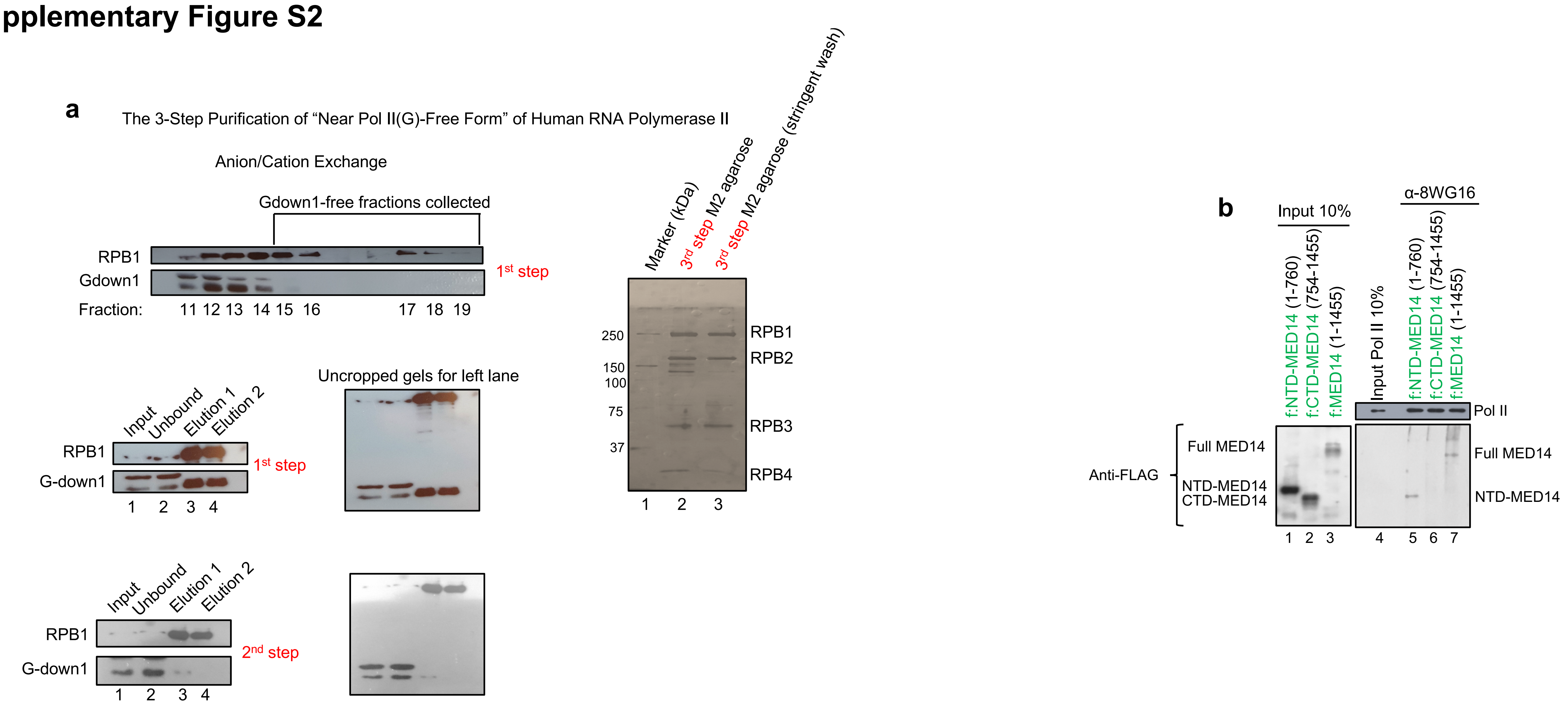

Recombinant FLAG-, His-, and HA-Tagged Pol II Subunits

### Supplementary Figure S3

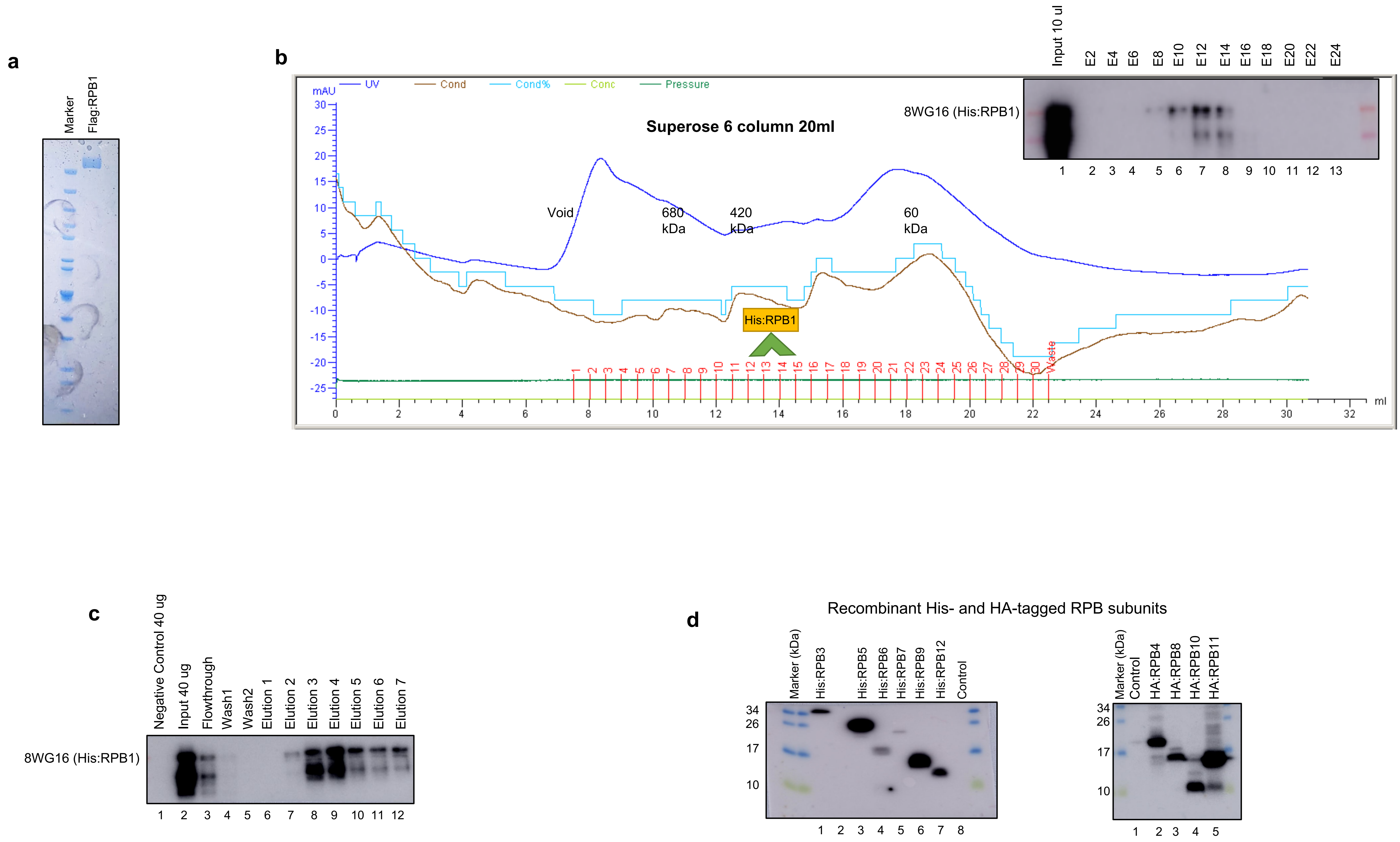

Characterization of MED14-NTD Human Core Interaction with Recombinant Pol II Subunits

### Supplementary Figure S4

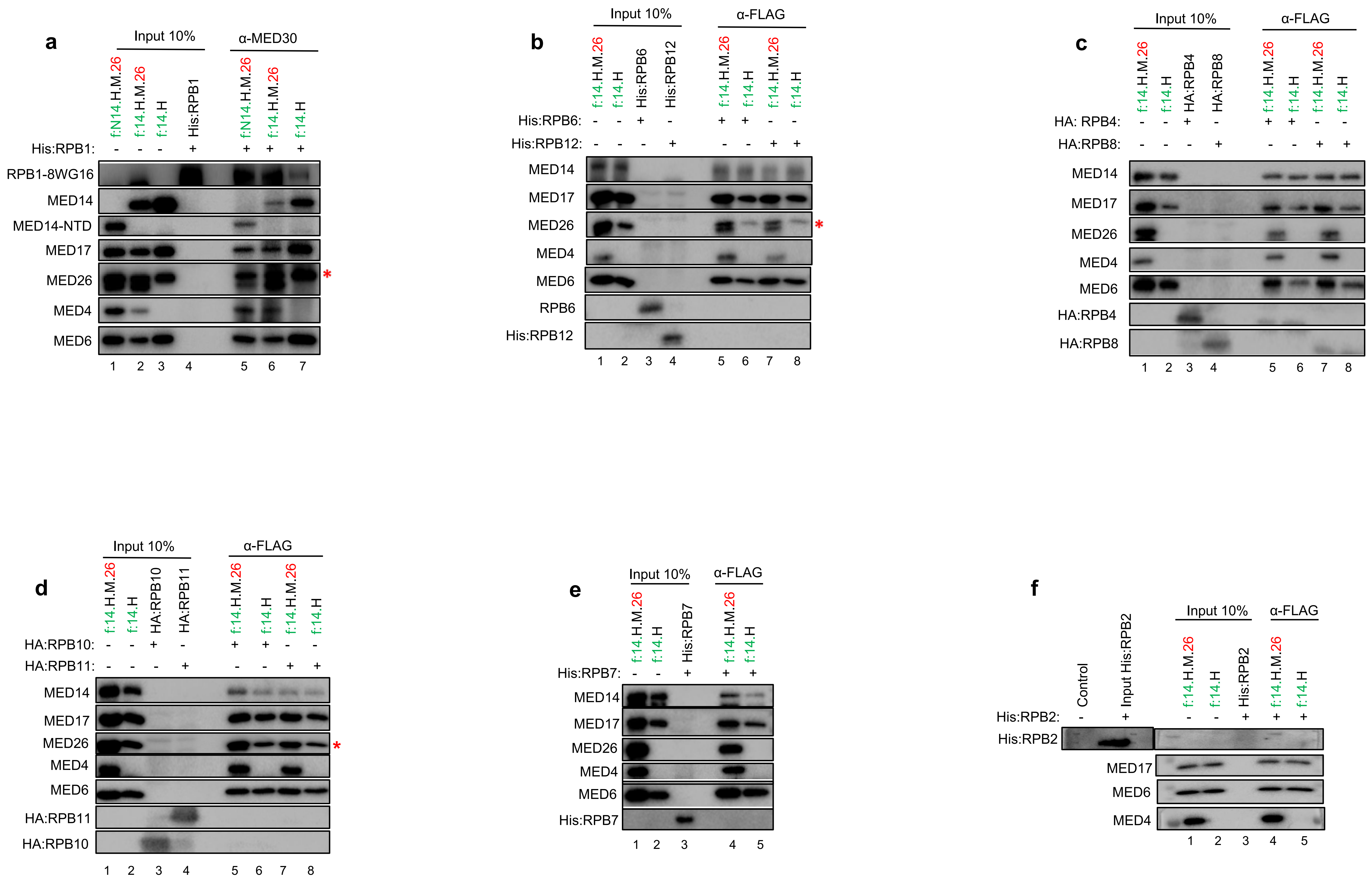

NTD-MED14 is Critical for Pol II Interaction

### Supplementary Figure S5

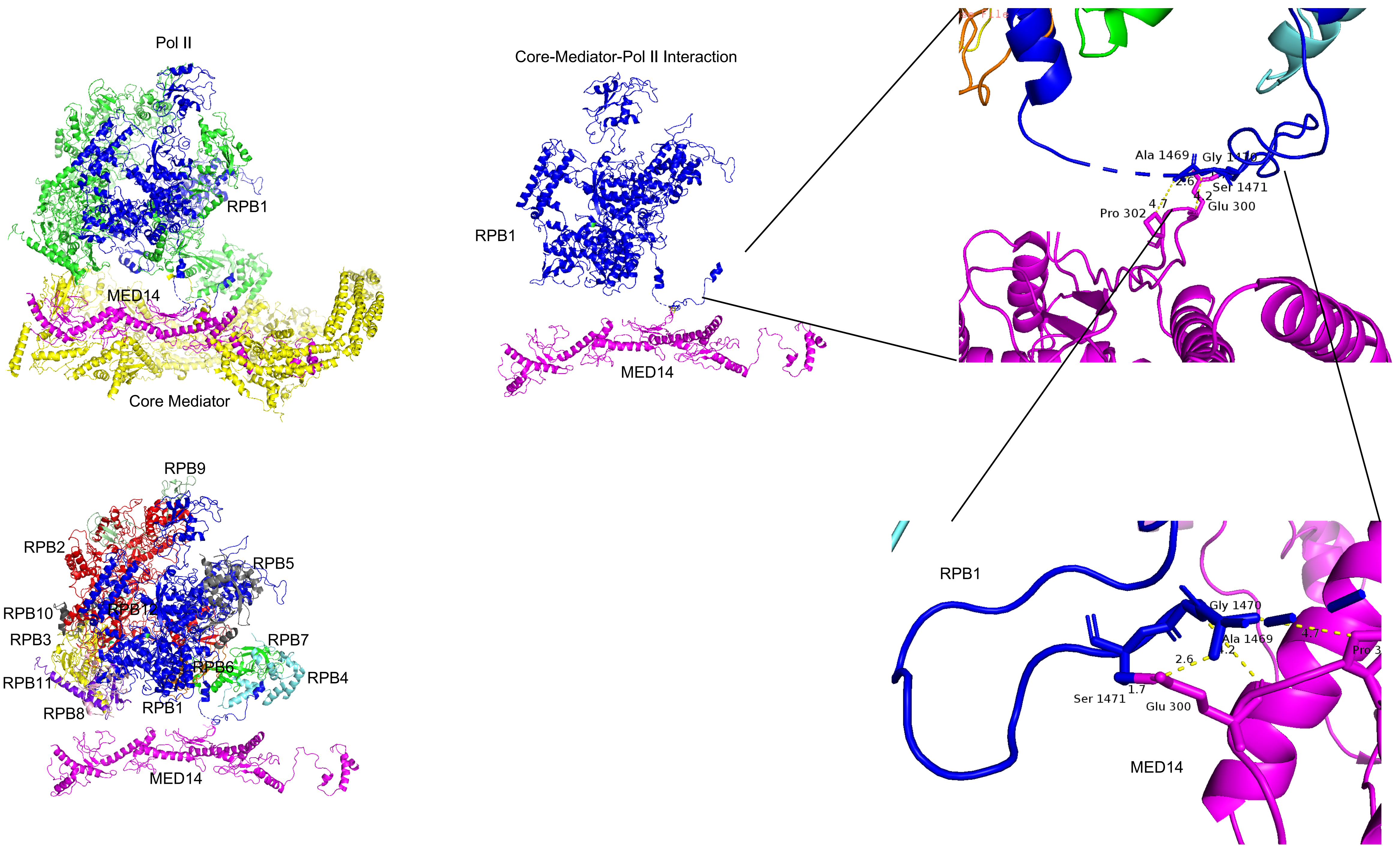
