## Supplementary Table 1 for "Mediator-RNA Polymerase II Interactions Critical for Transcriptional Activation Are Mediated by the N-terminal Half of MED14 and C-terminal Domain of RPB1"

### Protein ID: Q_9079277 *S. pombe* MED14

```
AA_QUERY      1   MEPPAIPHITEGFYPLPEIVETFSHHVLQELVSLAEVLPSMSNVEKKKKILDWLLRSRAFTMRLLVLARWVHLSPSVHRCIDVVAFLQGQ   90
SS_PSIPRED              HHHHHHHHHHHHHHHHHHHHHHHHHH  HHHHHHHHHHHHHHHHHHHHHHHHHHHHHHHHH HHHHHHHHHHHHHHHHHHH
SS_SPIDER2             EEHHHHHHHHHHHHHHHHHHHHHHHH   HHHHHHHHHHHHHHHHHHHHHHHHHHHHHHHH HHHHHHHHHHHHHHHHHH
SS_PSSPRED              HHHHHHHHHHHHHHHHHHHHHHHHHHHH HHHHHHHHHHHHHHHHHHHHHHHHHHHHHHHH  HHHHHHHHHHHHHH
SS_DEEPCNF              HHHHHHHHHHHHHHHHHHHHHHH   HHHHHHHHHHHHHHHHHHHHHHHHHHHHHHH  HHHHHHHHHHHHHH
CC_COILS_W28
CC_PCOILS_W28                                         CCCC
DO_DISOPRED3      DDD
DO_SPOTD          DD
DO_IUPRED

AA_QUERY      91  KFCFQNLVHVLQDIRYQLSFARLRNSDLVTALDILSTGTSLRLANAPTSKLYMLSESPLSTKQILQTLHALNMLIRIRLSLYEIIPTPFQ   180
SS_PSIPRED        HHHHHHHHHHHHHHHHHHH HHH   HHHHHHHHH            HHHHHHHHHHHHHHHHHHHHH     HHHH
SS_SPIDER2        HHHHHHHHHHHHHHHHHHHHHH       HHHHHH  E         HHHHHHHHHHHHHHHHHHHHH     H HE
SS_PSSPRED        HHHHHHHHHHHHHHHHHHHHHHHHHHH HHHHHHHHHHH    EEEE   EEEEE  HHHHHHHHHHHHHHHHHHHHHHHHHHH
SS_DEEPCNF         HHHHHHHHHHHHHHH        EEEEEE      EEEE  HHHHHHHHHHHHHHHHHHHHHHH      E
CC_COILS_W28        CCCCCCCCCCCCCCCCCCCCCCCCCCCCC
CC_PCOILS_W28     CCCCCCCCCCCCCCCCCCCCCCCCCCCCCCCCCCCCCCC
DO_DISOPRED3
DO_SPOTD
DO_IUPRED

AA_QUERY      181 HFTIANGRCTFTVPNEFSVSLTTNSQDPKSTGISFQWIVVDFQFHLPDFSSTPAKYRVFIELHLNEEIAAAFVLQKPILPLIYNILHKFC   270
SS_PSIPRED         EEE  EEEE  EEEEEEE     EEEEEEEEE   HHHHHHHHHHHHHHHHHHHHHHH   HHHHHHHHHHHHHH
SS_SPIDER2        EEEEE EEEEEE EEEEEEEE     EEEEEEEEEE   HHHHHHHHHHHHHHHHHHHHHH    HHHHHHHHHHHH
SS_PSSPRED         EEEE EEEEEE EEEEEEE    EEEEEEEEEEEE   EEEEEEEEE HHHHHHHHHH HHHHHHHHHHHHHHH
SS_DEEPCNF        EEEEE EEEEEE EEEEEEEE    EEEEEEEEEEEEE   EEEEEEEEEE HHHHHHHHH   HHHHHHHHHHHH
CC_COILS_W28
CC_PCOILS_W28
DO_DISOPRED3
DO_SPOTD
DO_IUPRED

AA_QUERY      271 LYQRLNLLSQQTFQLSRESWLGHLRGVYDEKPPRLRLYYWPQLNVKKEGKPGKIGHYIHIFVNTQPISAFERTLSSKRSSCEYDHFLLLV   360
SS_PSIPRED        HHHHHHHHHHHHHHHHHHH   EEEEEE  EEEEEE       EEEEEEE  HHHH       EEEE
SS_SPIDER2        HHHHHHHHHHHHHHHHHHH    EEEEE  EEEEEE      EEEEEEE  HHHHH      EEEE
SS_PSSPRED        HHHHHHHHHHHHHHHHHHHHHHHHHHHH     EEEEE       EEEEEEE  HHHHHHHH     EEEEEE
SS_DEEPCNF        HHHHHHHHHHHHHHHHHHH        EEEEEEE      EEEEEEE  HHHHHH     EEEEEE
CC_COILS_W28
CC_PCOILS_W28
DO_DISOPRED3
DO_SPOTD                            DDDDDDDD           DDDDDDD
DO_IUPRED

AA_QUERY      361 EWHHDGIVEHVPLDDHMDAQHLLLLITQKHAQLILEQIRKELHPNIFSEHVGGGLKIHVFDNEIIVKVNSVTGRLVLSSSASPLSPPRHL   450
SS_PSIPRED        EEEE EE    HHHHHHHHHHHHHHHHHHHHHHHHHHH     EEEEEE EEEEEEE  EEEEE   HH
```

```
SS_SPIDER2              EE    EE       HHHHHHHHHHHHHHHHHHHHHHHHH   EE    EEEEEE  EEEEEEE   EEEEEE      HHH
SS_PSSPRED              EEE   EEE      HHHHHHHHHHHHHHHHHHHHHHHHH  HHHHH   EEEEEE  EEEEEEEEEE EEEEE      HHHH
SS_DEEPCNF              EEE  EEEEEE    HHHHHHHHHHHHHHHHHHHHHHHHH         EEEEE  EEEEEE   EEEE       HHH
CC_COILS_W28
CC_PCOILS_W28
DO_DISOPRED3
DO_SPOTD
DO_IUPRED
```

```
AA_QUERY         451 RAAEKNIALNTQPPAQILNRLYFFCIQTQLLEVAQCAELHAVQGYYSFPYLTFSKGKWRKDGDSLWVLAYNVESNSWSVRLLNAAGQTLY   540
SS_PSIPRED           HHHHHHH    HHHHHHHHHHHHHHHHHHHHHHHHH  EE     EEEE       EEEEEEEE   EEEEEEE   EEE
SS_SPIDER2           HHHHHHHH   HHHHHHHHHHHHHHHHHHHHHHHHH  EEEE    EE   H     EEEEEEEE  EEEEEEE   EEE
SS_PSSPRED           HHHHHHHE   HHHHHHHHHHHHHHHHHHHHHHHHHHHHHHHHHH     EEE       EEEEEEEEE   EEEHHHHHH  EEE
SS_DEEPCNF           HHH       HHHHHHHHHHHHHHHHHHHHHHH           EEEEEEEEEEE  EEEEEEEEE  EEEEEEEE   EEE
CC_COILS_W28
CC_PCOILS_W28
DO_DISOPRED3
DO_SPOTD
DO_IUPRED              D D
```

```
AA_QUERY         541 TQDVHTTKGTLSIESFSRLSYLLEVQILLFNVQTACQARGMPFEYLPIPPKALIEDDFTTYVQTGCLCIMMPSSNEDMLPVVFVRAHDGQ   630
SS_PSIPRED           EEEE      HHHHHHHHHHHHHHHHHHHHHHHHHHHHH              EEEEE    HH       EEEEEEE    E
SS_SPIDER2           EEE       HHHHHHHHHHHHHHHHHHHHHHHHHHHHH                     E E       EEEEEE    E
SS_PSSPRED               EEEEHHHHHHHHHHHHHHHHHHHHHHHHHHHHHHH        HHH    HHHHHHH EEEEE        EEEEEEE
SS_DEEPCNF           EEE       HHHHHHHHHHHHHHHHHHHHHHHHHHHHH              HHHHHH  EEEE      EEEEEEEEE   E
CC_COILS_W28
CC_PCOILS_W28
DO_DISOPRED3
DO_SPOTD
DO_IUPRED
```

```
AA_QUERY         631 LIFDSRIKGKLPYQSETETEKNCYIDWRTGRITIRVQNFSSFEKTWIGLLKLVALSKTSAFNVDCITLKHVDFTYLDDEKFRATIHDDNT   720
SS_PSIPRED           EEEEEEEEE          EEEE   EEEEEE  HHHHHHHHHHHHHHHHHHHHH   EEEEEE  EEEEEE    EEEEEE
SS_SPIDER2           EEEEEEEE E          EEE   EEEEEE  HHHHHHHHHHHHHHHHHHHHH  E EEE   EEEEEE    EEEEEE
SS_PSSPRED           EEEE              EEEEEE  EEEEEEE   HHHHHHHHHHHHHHHH    EEEEEEE        HHHEEEE
SS_DEEPCNF           EEEEEEEEEE         EEEE   EEEEEE    HHHHHHHHHHHHHHEE     EEEEEEEEEEEE     EEEEEE  E
CC_COILS_W28
CC_PCOILS_W28
DO_DISOPRED3
DO_SPOTD
DO_IUPRED
```

```
AA_QUERY         721 FTLHFFNRHSPFHLISQFLQDTFSDGPSAIQPLRVIMDRTRGVLVAQELGYVVLARSLRQYRIILSKNHGIQVLLNRHGCILQDLSYLSA   810
SS_PSIPRED            EEEE     HHHHHHHHHHHHH  HHHHHHHHHHHHHHHHHHHHH   EEEEEE   EEEEEE   EEEEEEEEE   EEEEEEEE
SS_SPIDER2           EEEEE      HHHHHHHHHHHHH  HHHHHHHHHHHHHHHHHHHHH   EEEEEEE   EEEEEE    EEEEEEE    EEEEEEEEE
SS_PSSPRED           EEEEEE    HHHHHHHHHHHHH    HH HHHHHHHH   EEEHHHH HHHHHHHHHHHHHHHHH     EEEEHHH HHHHHHHHH
SS_DEEPCNF           EEEEEE   HHHHHHHHHHHHHH      HHHHHHHH         EEEEE    EEEEEE   HHHHHH    EEEEEEEE
CC_COILS_W28
CC_PCOILS_W28
```

```
DO_DISOPRED3
DO_SPOTD
DO_IUPRED
```

```
AA_QUERY      811  DSRYLEGTQTLTSQWEPCSWLNTVWEGDLGDDELNGQIEAAPEMHLIKMNKTADLTAILKRILAISRKK  879
SS_PSIPRED                HHH EE     EEE  EEE  HH HHHHHHHHHHHH
SS_SPIDER2                HHHHHHHHH     E  HHHHHHH  HHHHHHHHH
SS_PSSPRED                 HHHHHH       HHHHEE  HHHHHHHHHHHHHHH
SS_DEEPCNF               EE          EE  EEE   HHHHHHHHHHH
CC_COILS_W28
CC_PCOILS_W28
DO_DISOPRED3                                DDDD DDD
DO_SPOTD                                       DD
DO_IUPRED
```

SS = Secondary Structure; H = Alpha-helix; E= Beta-strand; CC = Coiled coils; D, DO = Disordered Region

#### Protein ID: Q_7346414 MED14 Human

```
AA_QUERY      1    MAPVQLENHQLVPPGGGGGGSGGPPSAPAPPPPGAAVAAAAAAAASPGYRLSTLIEFLLHRAYSELMVLTDLLPRKSDVERKIEIVQFAS    90
SS_PSIPRED                                           HHHHHHHHHHHH   EEHHHHHHHHHHHHHHHHHHHHHHHH    HHHHHHHHHHHHH
SS_SPIDER2                                             HHHHHHH     EEHHHHHHHHHHHHHHHHHHHHHHHH     HHHHHHHHHHHH
SS_PSSPRED            EEE                             HHHHHHHHHHHH    HHHHHHHHHHHHHHHHHHHHHHHH     HHHHHHHHHHHH
SS_DEEPCNF                                           HHHHHHHHH       HHHHHHHHHHHHHHHHHHHHHH      HHHHHHHHHHHH
DO_DISOPRED3       DDDD DDDDDDDDDDDDDDDDDDDDDDDDDDDDDDDDDDDDDD
DO_SPOTD           DDDDDDDDDDDDDDDDDDDDDDDDDDDDDDDDDDDDDDDDDDDDD
DO_IUPRED             DDDDDDDDDDDDDDDDDDDDDDDDDDDDDDDDDDD
```

```
AA_QUERY      91   RTRQLFVRLLALVKWANNAGKVEKCAMISSFLDQQAILFVDTADRLASLARDALVHARLPSFAIPYAIDVLTTGSYPRLPTCIRDKIIPP    180
SS_PSIPRED         HHHHHHHHHHHHHHHHH  HHHHHHHHHHHHHHHHHHHHHHHHHHHHHHHHHHHHHH          EEE              HH
SS_SPIDER2         HHHHHHHHHHHHHHHH   HHHHHHHHHHHHHHHHHHHHHHHHHHHHHHHHHHHHHH      HHHHHHHHH        HHHH
SS_PSSPRED         HHHHHHHHHHHHHHHHHH   HHHHHHHHHHH HHHHHHHHHHHHHHHHHHHHHHHHH   HHHHHHEEEEE        HHH
SS_DEEPCNF         HHHHHHHHHHHHHHHHH    HHHHHHHHH    EEEEE HHHHHHHHHHHHHHHH       EEEEEE          HHH
DO_DISOPRED3
DO_SPOTD
DO_IUPRED
```

```
AA_QUERY      181  DPITKIEKQATLHQLNQILRHRLVTTDLPPQLANLTVANGRVKFRVEGEFEATLTVMGDDPDVPWRLLKLEILVEDKETGDGRALVHSMQ    270
SS_PSIPRED             HHHHHHHHHHHHHHHHHHHHH   HHHH  EEE  EEEEEE  EEEEEEEE       EEEEEEEEEEEE           HHH
SS_SPIDER2             HHHHHHHHHHHHHHHHHHH     HHHEEEEE   EEEEEE  EEEEEEEEE      EEEEEEEEEEE            HHH
SS_PSSPRED             HHHHHHHHHHHHHHHHHHHH    HHH  EEE   EEEEEEE EEEEEEEEE      HHHHHHHHHEE       HHEHHHH
SS_DEEPCNF             HHHH   HHHHHHHHHHHH           EEE  EEEEEEEEEEEEEEEEE         EEEEEEEEE       HHHHHHH
DO_DISOPRED3
DO_SPOTD
```

```
DO_IUPRED

AA_QUERY      271  ISFIHQLVQSRLFADEKPLQDMYNCLHSFCLSLQLEVLHSQTLMLIRERWGDLVQVERYHAGKCLSLSVWNQQVLGRKTGTASVHKVTIK    360
SS_PSIPRED         HHHHHHHHHHHH     HHHHHHHHHHHHHHHHHHHHHHHHHHHHHHHHHHH EEEEEE    EEEEEEEE            EEEEEEE
SS_SPIDER2         HHHHHHHHHH      HHHHHHHHHHHH  HHHHHHHHHHHHHHHHH      EEEEEE    EEEEEEEE            EEEEEEE
SS_PSSPRED         HHHHHHHHHHHHH    HHHHHHHHHHHHHHHHHHHHHHHHHHHHHHHHH   EEEEHHH   EEEEEEE HHH       EEEEEEEEE
SS_DEEPCNF         HHHHHHHHHH        HHHHHHHHHHHHHHHHHHHHHHHHHHHHHHHH   EEEEEEE   EEEEEEEE           EEEEEEEE
DO_DISOPRED3
DO_SPOTD
DO_IUPRED

AA_QUERY      361  IDENDVSKPLQIFHDPPLPASDSKLVERAMKIDHLSIEKLLIDSVHARAHQKLQELKAILRGFNANENSSIETALPALVVPILEPCGNSE    450
SS_PSIPRED         E        EEEEE         HHHHHH       HHHHHHHHHHHHHHHHHHHHHHHHHH       EE     EEEEEE
SS_SPIDER2         E         EEEE     H   HHHHHH       HHHHHHHHHHHHHHHHHHHHHHHHH                EEEE E      E
SS_PSSPRED         E        HHH          HHHHHHHHHH  HHHHHHHHHHHHHHHHHHHHHHHHHHHH       HHHHHHHHHHHHH       H
SS_DEEPCNF         E        EEEE         HHHHHHHH      HHHHHHHHHHHHHHHHHHHHHHHHH                EEEEEE
DO_DISOPRED3
DO_SPOTD
DO_IUPRED

AA_QUERY      451  CLHIFVDLHSGMFQLMLYGLDQATLDDMEKSVNDDMKRIIPWIQQLKFWLGQQRCKQSIKHLPTISSETLQLSNYSTHPIGNLSKNKLFI    540
SS_PSIPRED         EEEEEEE    EEEEEE    HHHHHHHHHHHH  HHHHHHHHHHHHHHHHHHHHHHHHHH   EEE                   EEEE
SS_SPIDER2         EEEEEE     EEEEEE    HHHHHHHHHHH   HHHHHHHHHHHHHHHHHHHHHHHHH    EEEEE                 EEEE
SS_PSSPRED         HHHHHHHHHHHHHHHHHH     HHHHHHHHHHHHHHHHHHHHHHHHHHHHHHHHHHHHHH        EE               HHHH
SS_DEEPCNF         EEEEEEE    EEEEEEE     HHHHHHHHHHHHHHHHHHHHHHHHHHHHHHHHHHHH           E               EEEE
DO_DISOPRED3
DO_SPOTD
DO_IUPRED

AA_QUERY      541  KLTRLPQYYIVVEMLEVPNKPTQLSYKYYFMSVNAADREDSPAMALLLQQFKENIQDLVFRTKTGKQTRTNAKRKLSDDPCPVESKKTKR    630
SS_PSIPRED         EE     EEEEEEEEE     EEEEEEEEEEEEE       HHHHHHHHHHHHHH   EEE
SS_SPIDER2         EE     EEEEEEE        EEEEEEEEEEEEE      HHHHHHHHHHHHHHHH  EE                            H
SS_PSSPRED         HHHH   EEEEEEEEE       EEEEEEEEEEE       HHHHHHHHHHHHHHHHHHHH         HHHH              HH
SS_DEEPCNF         EE     EEEEEEEEE      EEEEEEEEEEEE       HHHHHHHHHHHHHH    E
DO_DISOPRED3                                                                         DDDDDDDDDDDDDDDDDDDDDDDD
DO_SPOTD                                                                            DDDDDDDDDDDDDDDDDDDDDDDDD
DO_IUPRED                                                                       DDD    DDDDDDDDDDDDDDD

AA_QUERY      631  AGEMCAFNKVLAHFVAMCDTNMPFVGLRLELSNLEIPHQGVQVEGDGFSHAIRLLKIPPCKGITEETQKALDRSLLDCTFRLQGRNNRTW    720
SS_PSIPRED             HHHHHHHHHHHHHH     HHHHHHHHHH        EE    EEEEEEE          HHHHHHHHHHH EEEEEE     EEE
SS_SPIDER2            HHHHHHHHHHHHHHHH     HHH HHH       E EEE     EEEEEE          HHHHHHHHH EEEEEEE       EE
SS_PSSPRED         HHHHHHHHHHHHHHHHHH        EEEEE         EEEE   HHHHHHHH         HHHHHHHHHHHHHHEEEE     HHH
SS_DEEPCNF                HHHHHHHHHHH     EEEEEEEE         EEE     EEEEEEEE        HHHHHHHHH     EEEEE     EE
DO_DISOPRED3       DD
```

```
DO_SPOTD          D
DO_IUPRED

AA_QUERY     721  VAELVFANCPLNGTSTREQGPSRHVYLTYENLLSEPVGGRKVVEMFLNDWNSIARLYECVLEFARSLPDIPAHLNIFSEVRVYNYRKLIL    810
SS_PSIPRED        EEEEEEE              EEEEEEEEE          HHHHHHHHHHHHHHHHHHHHHHHHHH           EEEEEEE  EEEE
SS_SPIDER2        EEEEEEE               EEEEEEEEEE         HHHHHHHHHHHHHHHHHHHHHHHHH       H  EEEEEEEE  EEEE
SS_PSSPRED        HHHHHHH                EEEEEHHHHH      HHHHHHHHHHHHHHHHHHHHHHHHHHH     HHHHHH EEEHHHHHHHHH
SS_DEEPCNF        EEEEEEE                EEEEEE           EEEEEE   HHHHHHHHHHHHHHHHH            EEEE    EEEE
DO_DISOPRED3
DO_SPOTD
DO_IUPRED

AA_QUERY     811  CYGTTKGSSISIQWNSIHQKFHISLGTVGPNSGCSNCHNTILHQLQEMFNKTPNVVQLLQVLFDTQAPLNAINKLPTVPMLGLTQRTNTA    900
SS_PSIPRED        EEE     EEEEEEE    EEEEEE            HHHHHHHHHHHH     HHHHHHHHHH HHHHHHHH
SS_SPIDER2        EEE    EEEEEEEE    EEEEEEEE          HHHHHHHHHHHH     HHHHHHHHHH HHHHHHH
SS_PSSPRED        HH      EEEEEE HH HEEEEEE           HHHHHHHHHHHHHH   HHHHHHHHHHH    HHHHH       E       HH
SS_DEEPCNF        EEE     EEEEEE     EEEEEE           HHHHHHHHHHHHHH    HHHHHHHHHH    HHHH
DO_DISOPRED3
DO_SPOTD
DO_IUPRED

AA_QUERY     901  YQCFSILPQSSTHIRLAFRNMYCIDIYCRSRGVVAIRDGAYSLFDNSKLVEGFYPAPGLKTFLNMFVDSNQDARRRSVNEDDNPPSPIGG    990
SS_PSIPRED          EEEE      EEEEEE  EEEEEEEEE   EEEEE                     HHHHHHHH    HHHHH
SS_SPIDER2         EEEEEEEE   EEEEEE  EEEEEEEEE   EEEEEE                E   HHHHHHHHHHHHHHHHH
SS_PSSPRED        HHHHH       HHHHHHHHHHEEEEEEE   EEEEE                     HHHHHHHHH    HHHH
SS_DEEPCNF         EEEEE      EEEEEE    EEEEEE    EEEEE                    HHHHHHHHH    HHHHH
DO_DISOPRED3                                                                               DDDDDDDDDDDD
DO_SPOTD                                                                                    DDDDDDDDDDDDDDDD
DO_IUPRED                                                                         D    DDDDDDDDDDDDDDDDDDDDD

AA_QUERY     991  DMMDSLISQLQPPPQQQPFPKQPGTSGAYPLTSPPTSYHSTVNQSPSMMHTQSPGNLHAASSPSGALRAPSPASFVPTPPPSSHGISIGP    1080
SS_PSIPRED
SS_SPIDER2            H
SS_PSSPRED        HHHHHHHHH
SS_DEEPCNF         HHHHHHH
DO_DISOPRED3             D  DDDDDDDDDDDDDDDDDDDDDDDDDDDDDDDDDDDDDD   DDDDDDDDDDDDDDDDDDDDDDDDDDDD   D
DO_SPOTD          DDDDDDDDDDDDDDDDDDDDDDDDDDDDDDDDDDDDDDDDDDDDDDDDDDDDDDDDDDDDDDDDDDDDDDDDDDDDDDDDDDDDDDDDDDDD
DO_IUPRED         DDDDDDDDDDDDDDDDDDDDDDDDDDDDDDDDDDDDDDDDDDDDDDDDDDDDDDDDDDDDDDDDDDDDDDDDDDDDDDDDDDDDDDDDDDDD

AA_QUERY     1081 GASFASPHGTLDPSSPYTMVSPSGRAGNWPGSPQVSGPSPAARMPGMSPANPSLHSPVPDASHSPRAGTSSQTMPTNMPPPRKLPQRSWA    1170
SS_PSIPRED
SS_SPIDER2
SS_PSSPRED                         EE                                                                    HHH
SS_DEEPCNF
```

```
DO_DISOPRED3                                        DDDDDDDDDDDDDDDDDDDDDDDDDDDDDDDDDDDDDDDDDDDDDDDDDDDDDDDD
DO_SPOTD                  DDDDDDDDDDDDDDDDDDDDDDDDDDDDDDDDDDDDDDDDDDDDDDDDDDDDDDDDDDDDDDDDDDDDDDDDDDDDDDDDDDD
DO_IUPRED                 DDDDDDDDDDDDDDDDDDDDDDDDDDDDDDDDDDDDDDDDDDDDDDDDDDDDDDDDDDDDDDDDDDDDDDDDDDDDDDDDDDDDDDDDDD
```

```
AA_QUERY     1171 ASIPTILTHSALNILLLPSPTPGLVPGLAGSYLCSPLERFLGSVIMRRHLQRIIQQETLQLINSNEPGVIMFKTDALKCRVALSPKTNQT    1260
SS_PSIPRED                HHHHHHHH                   HHHHHHHHHHHHHHHHHHHHH   EE       EEEEE  EEEEEEEE      E
SS_SPIDER2                HHHHHH                     HHHHHHHHHHHHHHHHHHHH HHHHH       EEEEE   EEEEEEE     EE
SS_PSSPRED        HHHHHHHHHHHHHHEEE           H HHHHHHHHHHHHHHHHHHHHHHHHHHHHHHHH     EEEEEE   EEEEE       E
SS_DEEPCNF             HHHHHHHHHH                    HHHHHHHHHHHHHHHHHHHHHHHH        EEEEEEE   EEEEEE      E
DO_DISOPRED3
DO_SPOTD
DO_IUPRED
```

```
AA_QUERY     1261 LQLKVTPENAGQWKPDELQVLEKFFETRVAGPPFKANTLIAFTKLLGAPTHILRDCVHIMKLELFPDQATQLKWNVQFCLTIPPSAPPIA    1350
SS_PSIPRED        EEEEE         HHHHHHHHHHHHHH       HHHHHHHHHHH   HHHHHHHHHHH HH         EEEEEEEEE
SS_SPIDER2        EEEEE         HHHHHHHHHHHHHHH      HHHHHHHHHHH   HHHHHHHHHHHHHH         EEEEEEEEEE
SS_PSSPRED        EEEEE          HHHHHHHHHHHHH       HHHHHHHHHHH  HHHHHHHHHHHHHHHH  HH HHHHEEEEEEEE
SS_DEEPCNF        EEEEE          HHHHHHHHHHH         HHHHHHHHHHH    EEE   EEEEEEEE        EEEEEEEEEE
DO_DISOPRED3
DO_SPOTD
DO_IUPRED            DDD
```

```
AA_QUERY     1351 PPGTPAVVLKSKMLFFLQLTQKTSVPPQEPVSIIVPIIYDMASGTTQQADIPRQQNSSVAAPMMVSNILKRFAEMNPPRQGECTIFAAVR    1440
SS_PSIPRED             EEE    EEEEEEEE           EEEEEEEEE     EEE           HHHHHHHHHHHHHHHH         HHHHHH
SS_SPIDER2             EEEE   EEEEEEEEE          EEEEEEEEE    EEEE           HHHHHHHHHHHHHHHH         HHHHHH
SS_PSSPRED             EEEEHHHHHHHHHHHH          EEEHHHHHH                  HHHHHHHHHHHHHHHHH       EHHHHHHH
SS_DEEPCNF             EEEE   EEEEEEEE           EEEEEEEEE                    HHHHHHHHHHHHHH          HHHHHH
DO_DISOPRED3                              DD D                  DDDDDDDDDDDDDDDD
DO_SPOTD
DO_IUPRED         D                                               DDDDDDDD
```

```
AA_QUERY     1441 DLMANLTLPPGGRP    1454
SS_PSIPRED        HHHHH
SS_SPIDER2        HHHHH
SS_PSSPRED        HHHHH
SS_DEEPCNF        HHHH
DO_DISOPRED3              DDDDDD
DO_SPOTD                    DDDD
DO_IUPRED               DDDDDDDD
```

SS = Secondary Structure; H = Alpha-helix; E= Beta-strand; CC = Coiled coils; D, DO = Disordered Region
